# Recognizing EEG responses to active TMS vs. sham stimulations in different TMS-EEG datasets: a machine learning approach

**DOI:** 10.1101/2025.06.29.662189

**Authors:** Ahmadreza Keihani, Francesco L. Donati, Simone Russo, Sara Parmigiani, Michela Solbiati, Adenauer G. Casali, Matteo Fecchio, Omeed Chaichian, John Rothwell, Marcello Massimini, Lorenzo Rocchi, Mario Rosanova, Fabio Ferrarelli

## Abstract

**Background:** Transcranial Magnetic Stimulation (TMS) with simultaneous Electroencephalogram (TMS-EEG) allows assessing the neurophysiological properties of cortical neurons. However, TMS-evoked EEG potentials (TEPs) can be affected by components unrelated to TMS direct neuronal activation. Accurate, automatic tools are therefore needed to establish the quality of TEPs.

**Objective:** To assess the discriminability of EEG responses to TMS vs. EEG responses to sham stimulations using sequence-to-sequence machine learning (ML).

**Methods:** Two indipendent TMS-EEG datasets including TMS and several sham stimulation conditions were obtained from the left motor area of healthy volunteers (N=33 across datasets). A Bi-directional Long Short-Term Memory (BiLSTM) ML network was used to label each time point of the EEG signals as pertaining to TMS or sham conditions. Main outcome measures included accuracy at single-trial level and after averaging five to twenty trials.

**Results:** For TMS conditions, post-stimulus vs. baseline/pre-stimulus EEG comparisons yielded moderate (60%-75%) single-trial accuracy and high-accuracy (>75%) for 20 trials across datasets, while for sham conditions post- vs. baseline/pre-stimulus EEG comparisons yielded lower accuracy rates than for TMS conditions, except for unmasked auditory stimulation. Furthermore, baseline/pre-stimulus TMS vs. baseline/pre-stimulus sham EEG comparisons showed chance-level accuracy, whereas post-stimulus TMS vs. post-stimulus sham EEG comparisons had moderate (single trial) to high (20 trial) accuracy, except for TMS with and without the click noise masking. Single-subject findings were comparable to group-level results across datasets.

**Conclusions:** TEPs after active TMS are discernible from various sham stimulations even after a handful of trials and at the single-subject level using a BiLSTM ML approach.

## Introduction

The combination of TMS with simultaneous EEG (TMS-EEG) provides a unique method to non-invasively, directly target cortical neurons, thus allowing for the investigation of the neurophysiological properties of different cortical areas [1–3]. EEG responses to single TMS pulses, commonly described as transcranial evoked potentials (TEPs), are characterized by early components, which are determined by local excitatory cortical pyramidal neurons and inhibitory interneurons [4, 5], and later components reflecting the interplay between cortical and subcortical regions [6–8]. However, TEPs can also be affected by several confounding factors. First, TMS produces a “click” upon discharge, which generates an auditory-evoked potential (AEP) through air and bone conduction [6]. Moreover, the TMS-induced electric field can activate craniofacial muscles and free cutaneous nerve endings in the scalp, thus producing somatosensory evoked potentials (SEPs) [9]. These confounds may hinder an accurate interpretation of TEPs. Although dedicated tools to help address those confounds have been recently introduced [10], experimental countermeasures like playing a TMS click masking noise [11] are not always utilized in TMS-EEG studies and have been employed with various degrees of effectiveness [12–14].

Recent TMS-EEG studies aiming at differentiating TEPs due to TMS direct cortical activation from sensory responses yielded contradictory findings. Gordon and colleagues [15] reported clear distinctions between TEPs and EEG responses to auditory/somatosensory stimulations, while Conde et al. [13] found no significant differences between sham and TMS, concluding that TEPs primarily arise from auditory and somatosensory activation. In contrast, two recent studies found that TMS, coupled with appropriate masking of sensory input, resulted in lateralized responses at the stimulation site lasting ∼300 ms, whereas sham stimulations yield sensory evoked responses, represented by late (100-200 ms) components, mostly located at central scalp regions [16, 17]. Based on these results [16, 17], it was concluded that TMS, when controlling for confounding sources, produces responses primarily from direct neuronal activation. Yet, there is a lack of data-driven approaches that can systematically identify EEG responses to TMS and sham stimulations [12, 18].

Machine learning (ML) algorithms have gained increasing attention in EEG research based on their ability to differentiate conditions [19–22]. These algorithms excel in uncovering hidden relationships within complex datasets, providing insights otherwise precluded. ML algorithms are also well-suited for handling large, complex datasets where the number of inputs surpasses the number of subjects, as for TMS-EEG data, which require differentiating temporal and spatial information. In these cases, traditional statistical approaches often yield suboptimal results [16]. Hence, the utilization of ML represents an intriguing yet underexplored approach in the TMS-EEG field.

One of those ML-based approaches involves entering all spatio-temporal information from TMS-EEG data in a recurrent neural network (RNN) algorithm [23–28], which can then assess whether each time point is originating from TMS or sham conditions. A recent study used ML-based classification and showed that it could disentangle TEPs from the TMS click related AEPs better than conventional statistical analyses [29]. However, the ML performance was assessed: 1) on EEG responses across all trials, rather than on responses from single/few trials; 2) on a single TMS-EEG dataset; 3) only on auditory stimulation as confound; 4) withouth accounting for baseline differences between TMS and sham sessions; and 5) at the group, but not at the single subject level, thus leaving several unanswered questions concerning TMS vs. sham responses [6, 29]. First, can we identify TEPs with at least moderate accuracy at the single trial level using ML? And can high accuracy be achieved after averaging a small number of trials? Second, can we leverage ML to identify TEPs with high accuracy in different datasets? Third, can ML identify TEPs more accurately than both somatosensory and auditory evoked potentials when comparing each TMS and sham condition is compared to its baseline? Fourth, can this ML approach reach high accuracy when comparing the post-TMS vs. the post-sham stimulation responses, but low-chance level when comparing their baselines? Finally, can high accuracy for TEPs vs. several sham responses be achieved at the single-subject level?

Here, we began addressing those questions by comparing TMS and several sham stimulation sessions in different TMS-EEG datasets using bidirectional sequence-to-sequence long-short term memory (BiLSTM) ML [23–28]. The BiLSTM assessed TMS-EEG data bidirectionally (both forward and backward in time), used sequence-to-sequence (labeled each time point as TMS or sham), analyzed single trial and 5-20 averaged trials, compared TMS and sham sessions at baseline (pre-TMS vs. pre-sham) and after stimulation (post-TMS and post-sham), and examined both group and single subject performance.

## Material and methods

### Datasets

The BiLSTM ML model was applied to two datasets collected on thirty-three healthy subjects (HS) by two different research groups [16, 17]. Dataset 1 (N=14 HS) included: a) The condition to optimize cortical activation by TMS (effective TMS, eTMS), i.e., TEPs recorded with the TMS coil touching the scalp and the TMS click masked by a custom-made noise (TMS1) [10]; b) Realistic sham, recorded with the TMS coil tilted 90 degrees, simultaneous electrical stimulation to produce a TMS-like scalp sensation, and the TMS masking noise (S1); c) AEP not masked, obtained with the coil at 90 degrees and no masking noise (AEP1); d) AEP masked, as in c, but with noise masking (AEPm1); and e) High intensity electrical stimulation, obtained with an electrical stimulation producing discomfort in all subjects (HES1). Dataset 2 (N=19 HS) included: a) TEP, recorded with the TMS coil on the scalp while playing a masking noise (TEP, TMS2); b) Sham, with a sham coil on the scalp and noise masking (S2); c) AEP not masked, with the TMS coil on a 5 cm pasteboard cylinder placed on the scalp, but without noise masking (AEP2); d) AEP masked, as in c but with noise masking (AEPm2); e) Low-intensity ES, with an electrical stimulation of the scalp of intensity similar to that generated by 90% resting motor threshold (RMT) TMS (LES2); and f) TEP not masked, with the coil on the EEG cap, but no noise masking (TMSnm2). Both datasets utilized a neuronavigation system to target the stimulated cortical area (i.e., the primary motor cortex, M1) [16, 17]. However, for Dataset 1 a visualization of TEPs (rt-TEP [30]), which allows optimizing TMS parameters (coil position, stimulation intensity) based on EEG responses, was employed [17], while for Dataset 2 a fixed TMS intensity (i.e., 90%RMT) with no rt-TEP was chosen [16]. Furthermore, Dataset 1 TMS-EEG data collected more trials (217.64±30.70) than Dataset 2 (106±4.56).

### Analytical method, machine-learning approach

A BiLSTM ML network, which retain information for longer periods of time and allows for capturing long-term dependencies, was employed [23–26, 31]. BiLSTM is well suited for TMS-EEG data because it leverages both forward and backward information during training and inference, which can be utilized to distinguish temporal patterns (Figure 1). We used Statistics and Machine Learning Toolbox in MATLAB 2022a and a BiLSTM architecture, wherein input to the network were samples from all EEG channels and a time window of 300 ms for both pre-stimulus [-500 ms, -200 ms] and post-stimulus [20 ms, 320 ms]. The BiLSTM layer included 5 hidden state units and was set to output sequence. To mitigate overfitting, a dropout layer with 0.4 probability rate was added after the BiLSTM layer. A fully connected layer was included, which mapped the BiLSTM layer output to the number of classes, which was set to 2 (e.g., TEP vs. sham). A softmax layer was added to generate the final classification output. The model used data from all EEG channels for the defined time windows. Forward and backward activations were used to calculate the output at each time point (i.e., *y_ti_*), which was labeled as sham or TEP as a binary sequence with the same length as the input sequence from each trial.

**Figure 1.**
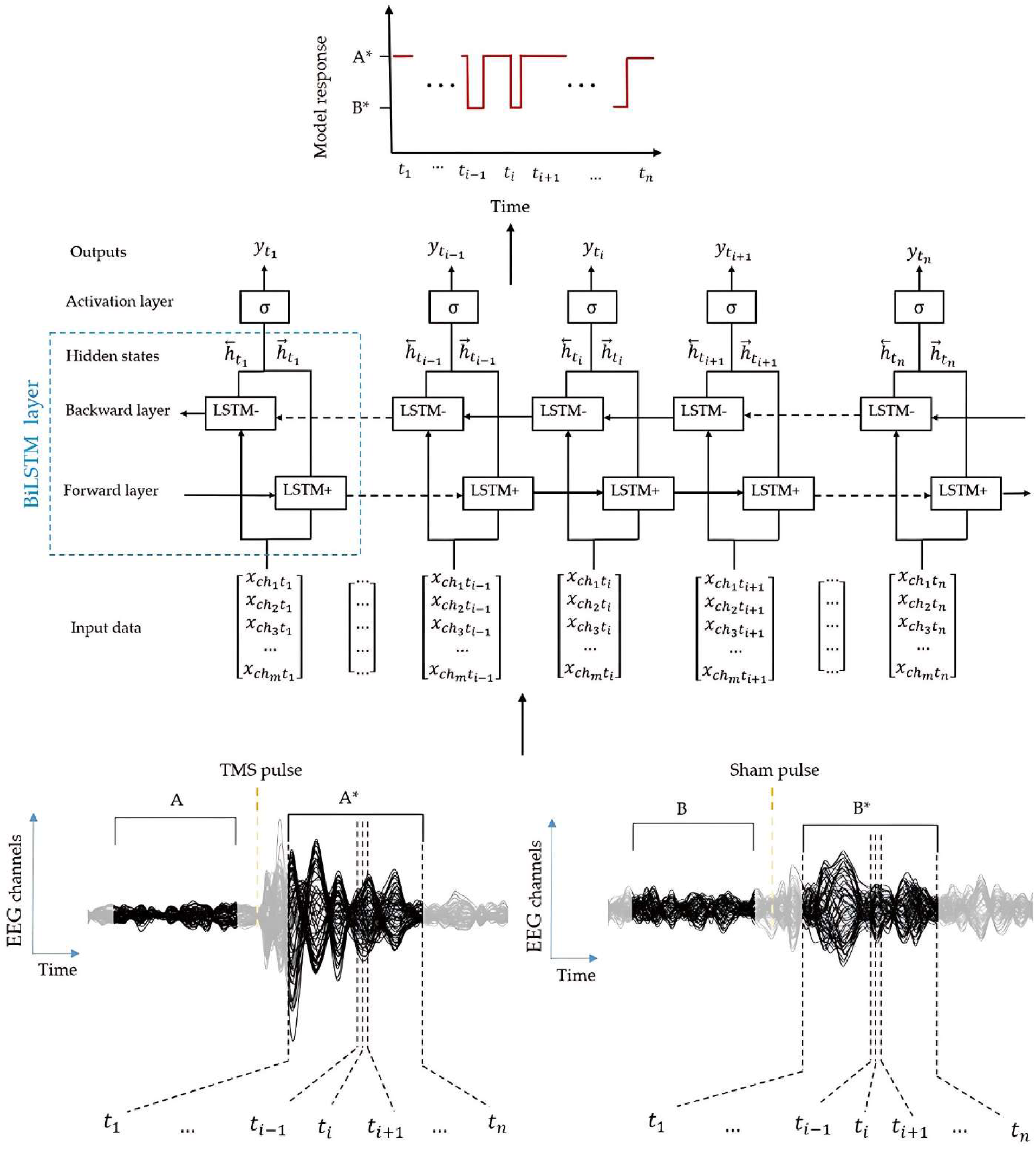
The bidirectional sequence-to-sequence long-short term memory (LSTM) machine-learning approach. At each time point t, the input multichannel time-series data are processed by a bidirectional LSTM layer. The resulting hidden states from the forward and backward LSTM layers are concatenated and passed through dropout and fully connected layers, followed by a softmax operation (collectively denoted as the ’activation layer’ for simplicity in the figure) to produce the output class probabilities. An example of our machine-learning approach applied to TMS/EEG sessions is shown.

The network was trained using the Adam optimizer [32] with a maximum of 60 epochs. The training progress was executed in a parallel computing environment for efficient processing. To assess the model’s performance, we employed leave one out (LOO), a subject-independent validation method that involves removing one subject from the training data and using such data for testing iteratively. To assess reliability of the LOO findings, a subject-dependent 5-fold cross-validation (CV) approach was utilized. For CV, trials from each dataset were divided into multiple folds (in every fold, 80% data [i.e., all trials of all subjects for sham and TEP] were used as a training set and 20% as a test set).

We first used LOO on the conditions shared across the two datasets (i.e., TEP [TMS1, TMS2], sham [S1, S2], AEP not masked [AEP1, AEP2], and AEP masked [AEPm1, AEPm2]) to assess the consistency of ML performance. For these conditions, we performed within-condition (i.e., post-stimulus vs. pre-stimulus) and between-condition (i.e., pre-stimulus vs. pre-stimulus and post-stimulus vs. post-stimulus) comparisons for both datasets. Within-condition comparisons established how well the ML algorithm differentiated post stimulus responses from their baselines, whereas between-condition comparisons examined whether BiLSTM could discern TMS1 and TMS2 from their respective sham conditions when looking at post stimulus (i.e., post-TMS vs. post-sham comparisons), but not when comparing their baselines (i.e., pre-TMS vs. pre-sham comparisons). We then applied the same approach to the conditions unique for Dataset 1 (HES1), and Dataset 2 (Low intensity ES [LES2] and TEP not masked [TMSnm2]). Accuracy (overall proportion of correct predictions), which was categorized as low (<60%), moderate (60%-75%), and high (>75%) [33], was the primary outcome measure for single trial and for 5-20 trials analyses. When averaging, we accounted for the number of trials (i.e., decreasing trials available for training and testing by averaging) and adjusted performance accordingly (see supplementary figures 2-19). Besides accuracy, sensitivity, specificity, area under the curve (AUC), and F1 score were computed. These parameters were also calculated at the single subject level. Due to differences in experimental set-up and sham conditions across datasets, BiLSTM was applied to each dataset separately. Thus, instead of training a model on one dataset and testing it on another, we examined how well our ML model distinguished TEP and sham conditions within each dataset.

Bottom panel) TMS/EEG traces from one representative subject from Dataset1. The model employs user-defined time windows (i.e., 300 ms long) relative to the TEP or sham pulse as input. In this example: A* (left traces, post TMS) vs B* (right traces, post sham). Of note, the model can also be employed to compare time windows within the same condition (e.g.: A vs A* for pre. vs. post TMS). Middle panel) general block diagram of Model; The model uses the entire spatio-temporal information (all channels [channel number (*Ch_i_*) at each time point [sample time (*t_i_*)]) as input data, while forward and backward directions are shown as black arrows and h represent the hidden states of the model. The forward and backward layers of the model are displayed as LSTM+ and LSTM-, respectively. Top panel) model output; in this example, the output of the model consists of a binary sequence, which defines whether the response label at each time point is TMS (A*) or sham (B*)

## Results

### Within-condition (post-stimulus vs. pre-stimulus for TMS and sham) comparisons

#### TMS1 and TMS2 had single trial moderate accuracy and high accuracy with 20 trials

In Dataset 1, the post vs. pre TMS comparison (TMS1) yielded moderate accuracy for single trial and high accuracy for 20 trials (Figure 2, top left panel, blue solid line, and Table 1), Acc range [mean Acc of single trial-mean Acc of 20 trials averaged across subjects]=[71.23%-100%]. Similarly, in Dataset 2 the post vs. pre TMS comparison (TMS2) showed moderate to high accuracy from single trial to 20 trials (Figure 2, top right panel, blue solid line, and Table 1, Acc range=[61.32%-77.62%]). However, accuracy levels for TMS2 were lower than for TMS1 (Figure 2, Table 1). This finding can be explained by a higher signal to noise ratio (SNR) for TMS1 relative to TMS2 [TMS1 SNR=4.24±0.56, TMS2 SNR=2.96±1.56, tstat=2.35, pval=0.025], likely due to using the rt-TEP to optimize TMS doses for TMS1 vs. fixed TMS intensities for TMS2. We also performed surrogate data analysis (i.e., artificial TEPs created through random circular shift of the data) for TMS1 and TMS2, which yielded chance level accuracy for both datasets (Supp. Figure 1), further suggesting that accuracy differences between TMS1 and TMS2 were related to TEPs. Combined, these findings indicate that TEPs, especially when collected with high SNR, can be detected even after a handful of TMS-EEG trials.

**Figure 2.**
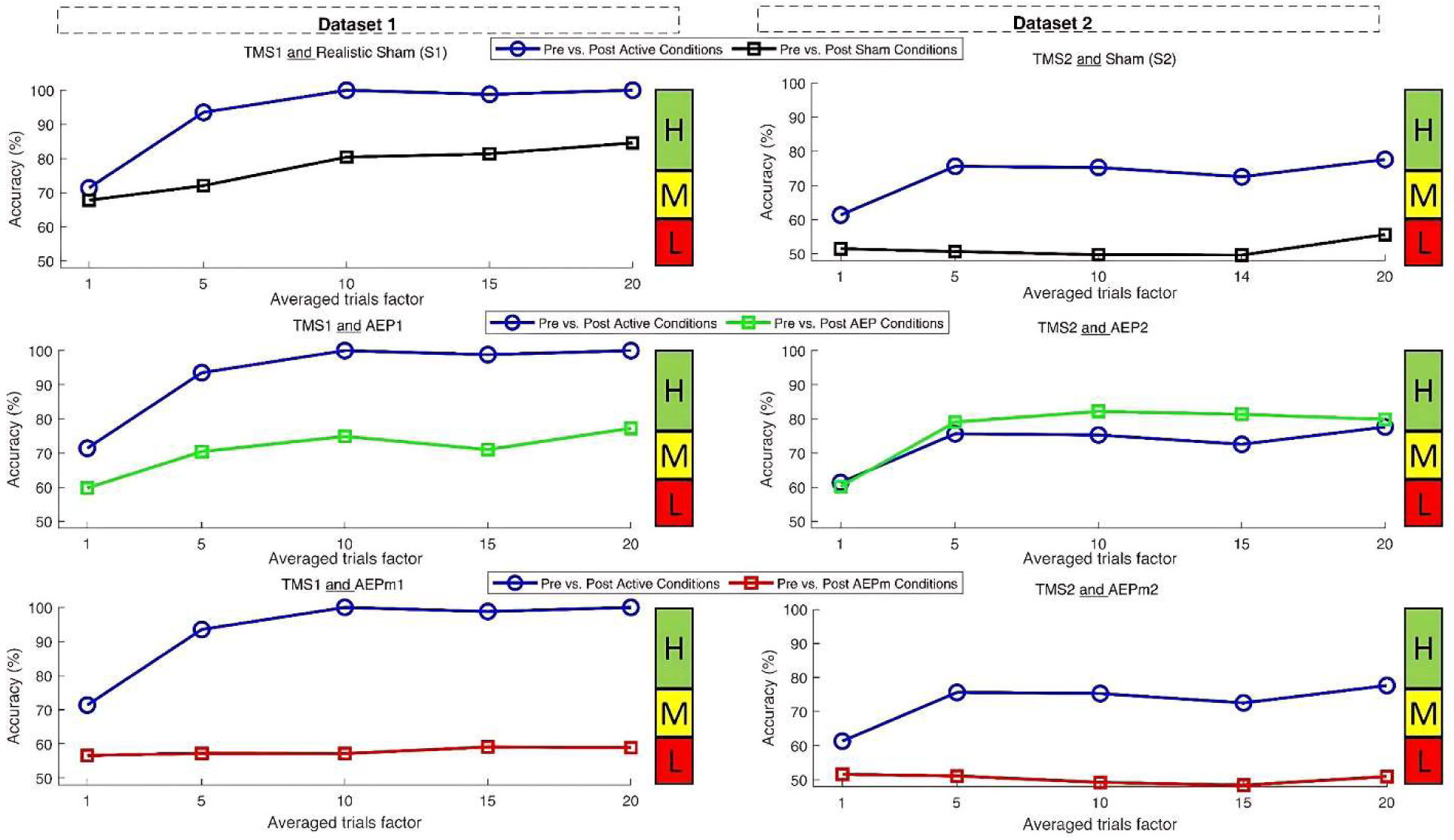
TMS1 and TMS2 had moderate (single trial) to high (average of 20 trials) accuracy for within-condition, post- vs. pre- comparisons that was higher than the accuracy of the shared sham conditions, expect AEP2, across datasets. Accuracy rates (%, y axes) after averaging an increasing number of trials (N, x axes) for post- vs. pre- stimulus comparisons of TEP (blue traces) and sham conditions (black, green, and red traces) shared across datasets. Color bar chart shows Acc ranges: Low (L < 60%), Medium (60% ≤ M ≤ 75 %), and High (H > 75 %).

**Table 1:**
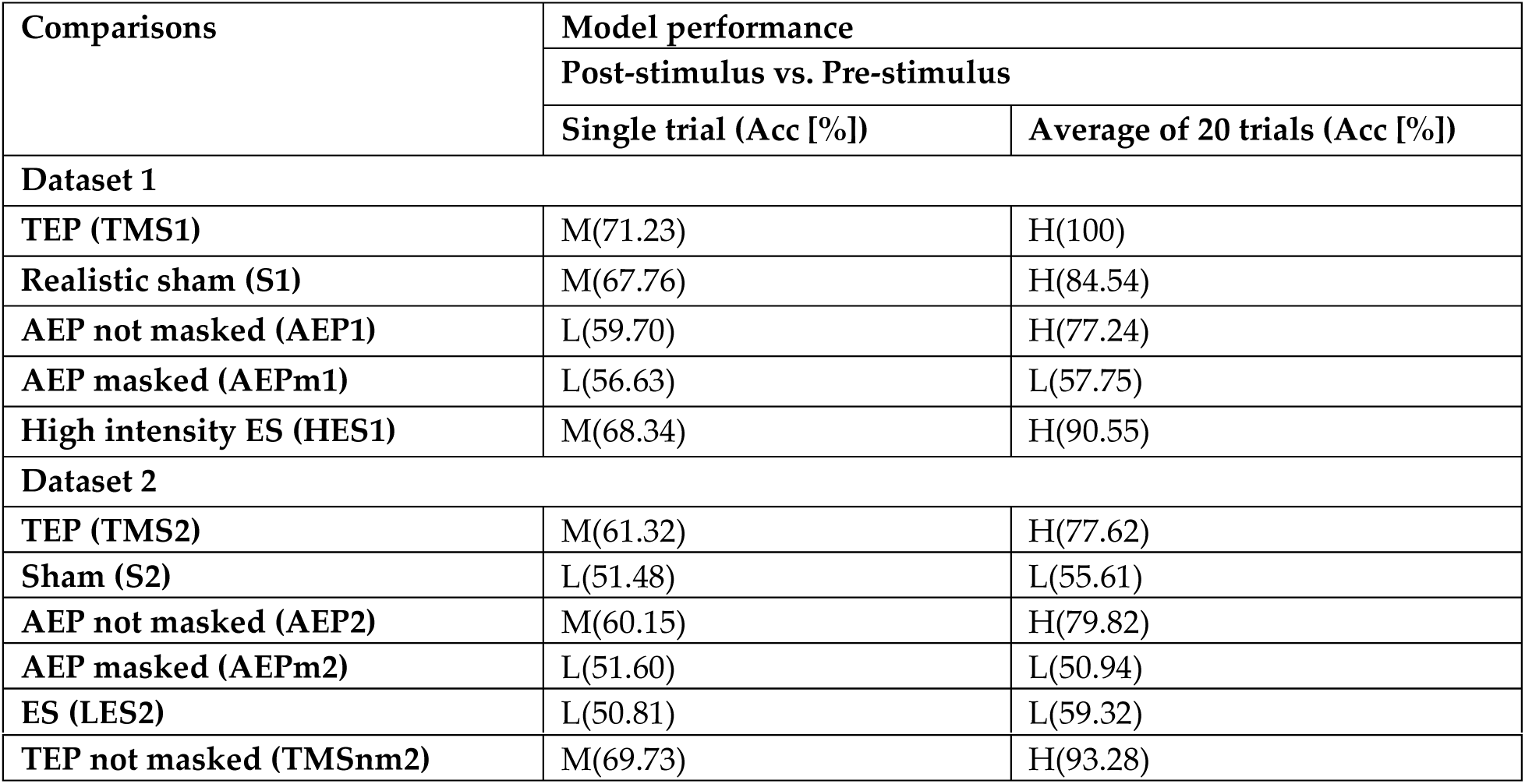
Main findings from within-condition comparisons of TEP and sham stimulations. L: low accuracy (<60%), M: moderate accuracy (60-75%), H: high accuracy (>75%) for single trial Acc vs. average of 20 trials Acc across subjects.

| Comparisons | Model performance |  |
| --- | --- | --- |
|  | Post-stimulus vs. Pre-stimulus |  |
|  | Single trial (Acc [%]) | Average of 20 trials (Acc [%]) |
| **Dataset 1** |  |  |
| TEP (TMS1) | M(71.23) | H(100) |
| Realistic sham (S1) | M(67.76) | H(84.54) |
| AEP not masked (AEP1) | L(59.70) | H(77.24) |
| AEP masked (AEPm1) | L(56.63) | L(57.75) |
| High intensity ES (HES1) | M(68.34) | H(90.55) |
| **Dataset 2** |  |  |
| TEP (TMS2) | M(61.32) | H(77.62) |
| Sham (S2) | L(51.48) | L(55.61) |
| AEP not masked (AEP2) | M(60.15) | H(79.82) |
| AEP masked (AEPm2) | L(51.60) | L(50.94) |
| ES (LES2) | L(50.81) | L(59.32) |
| TEP not masked (TMSnm2) | M(69.73) | H(93.28) |

#### Sham conditions shared across datasets showed variable accuracy levels but, except for AEP2, had lower single and averaged trial accuracy relative to TMS conditions

For Dataset 1, realistic sham (S1) (Figure 2, top left panel, black solid line, and Table 1, Acc range=[67.76%-84.54%]) and AEP not-masked (AEP1) (Figure 2, middle left panel, green solid line, and Table 1, Acc range=[61.32%-77.62%]) had moderate to high accuracy from single trial to 20 trials, whereas AEP masked (AEPm1) showed low accuracy and negligible differences between single and averaged trials (Figure 2, lower left panel, red solid line, and Table 1, Acc range=[56.63%-57.75%]). Furthermore, S1, AEP1 and AEPm1 all had lower accuracy for both single and averaged trials relative to TMS1 (Table 1).

For Dataset 2, post vs. pre AEP2 showed moderate single trial accuracy that increased to high accuracy with averaging (Figure 2, middle right panel, green solid line, and Table 1, Acc range=[60.15%-79.82%]), whereas low, close-to-chance accuracy levels were obtained for AEPm2 (Acc range=[50.94%-51.50%], Figure 2, bottom right panel, red solid line and Table 1), and S2 in both single trial and 20 trials (Figure 2, top right panel, black solid line, and Table 1, Acc range=[51.48%-55.61%]). All those sham conditions, except AEP2, had lower accuracy for both single and averaged trial analyses relative to TMS2 (Table 1).

#### Unique sham conditions showed moderate to high accuracy for HES1 and TMSnm2 and chance-level accuracy for LES2 vs. their baselines

Regarding sham conditions unique to each dataset, in Dataset 1, HES1 showed moderate single trial accuracy that increased with averaging (accuracy range=[68.34%-90.55%], Supp. Figures 2, and Table 1), while in Dataset 2 post vs. pre comparison for LES2 showed chance-level single trial accuracy slightly increasing with averaging (accuracy range=[50.81%-59.32%], Supp. Figures 3, and Table 1). Both sham conditions had lower single and averaged trial accuracy compared to TMS. Dataset 2 also included TMS not masked (TMSnm2), which yielded moderate single trial accuracy that increased with averaging (accuracy range= [69.73%-93.28%], Supp. Figure 4 and Table 1). As expected, given that it included both TEP and AEP components, this sham condition reached higher accuracy than TMS2.

#### Single subject and other performance parameters confirmed group accuracy findings

For all these within-condition comparisons, single-subject analyses confirmed what was observed in group analyses and showed that group effects were not driven by just a few individuals but were present in most subjects (Figure 3, single trial in blue circles and 20 trials in green asterisk). Nonetheless, some variability was observed in individual accuracy for TEP and sham conditions across datasets.

**Figure 3.**
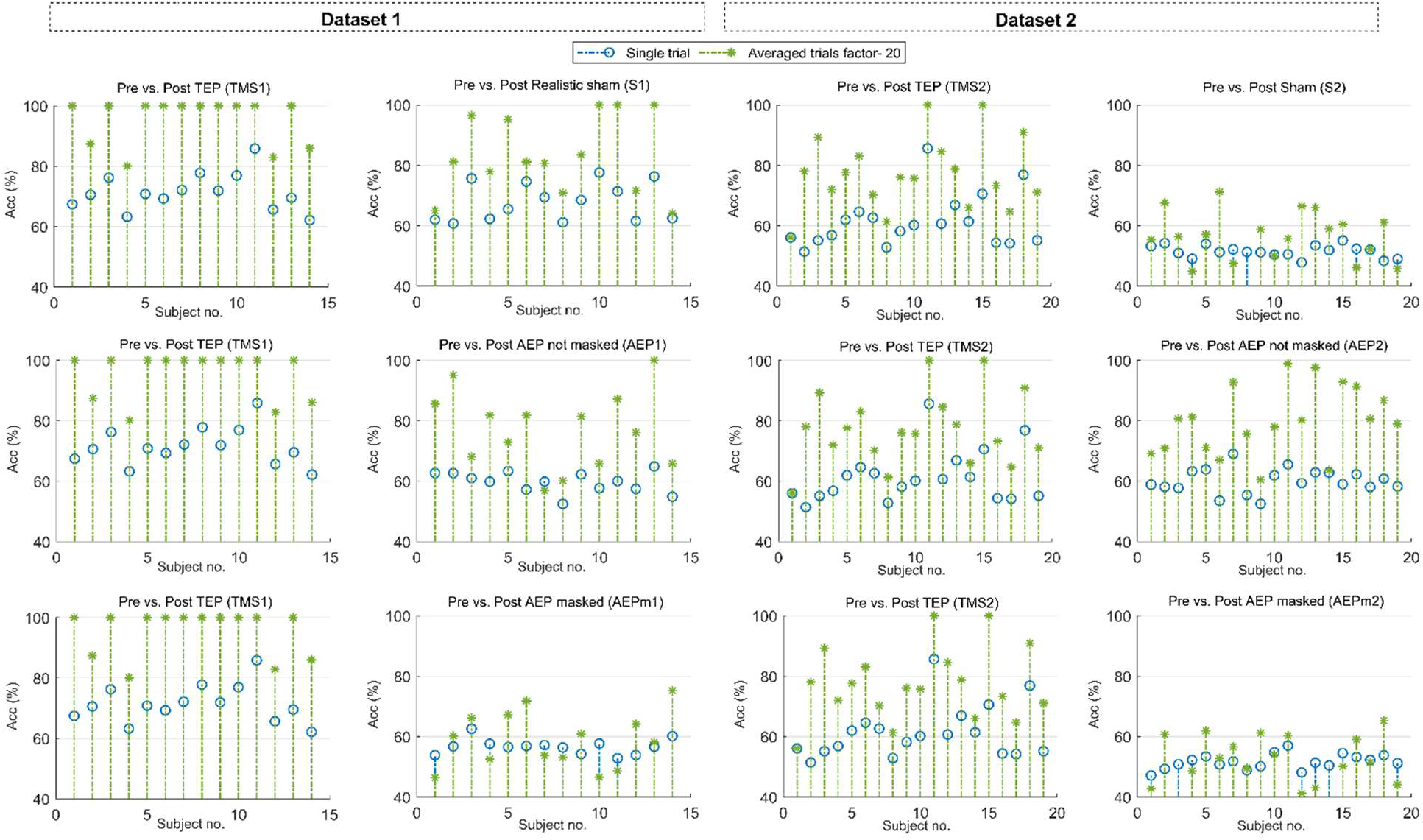
Single subject confirmed group-level findings for within-condition, post- vs. pre- comparisons of TMS and shared sham conditions across datasets. Individual accuracy rates for single trial (blue circles) and 20 trials (green asterisks) of TMS (first and third column) and shared sham conditions (second and fourth column) are shown for both datasets.

Sensitivity, specificity, AUC, and F1-score provided similar results (Supp. Figures 2-10) to those observed with accuracy for both datasets. Furthermore, CV yielded results in confirmation of LOO across datasets (see Supp. Figures 11-19).

### Between-condition (pre-TMS vs. pre-sham and post-TMS vs. post-sham) comparisons

#### Baseline/pre-stimulus comparisons of TMS vs. sham conditions shared across datasets resulted in low/chance level accuracy

In Dataset 1, baseline/pre-stimulus comparisons yielded low accuracy rates for single trial with marginal increases after 20 trials for TMS1 vs. S1 (Figure 4, top left panel, black dotted line, and Table 2, Acc range=[54.87%-55.49%]), TMS1 vs. AEP1 (Acc range=[55.62%-59.69%], Figure 4, middle right panel, green dotted line, and Table 2), and TMS1 vs. AEPm1 (Acc range=[57.41%-62.79%], Figure 4, bottom left panel, red dotted line, and Table 2). Dataset 2 showed similar results (Figure 4, right panels, and Table 2).

**Figure 4:**
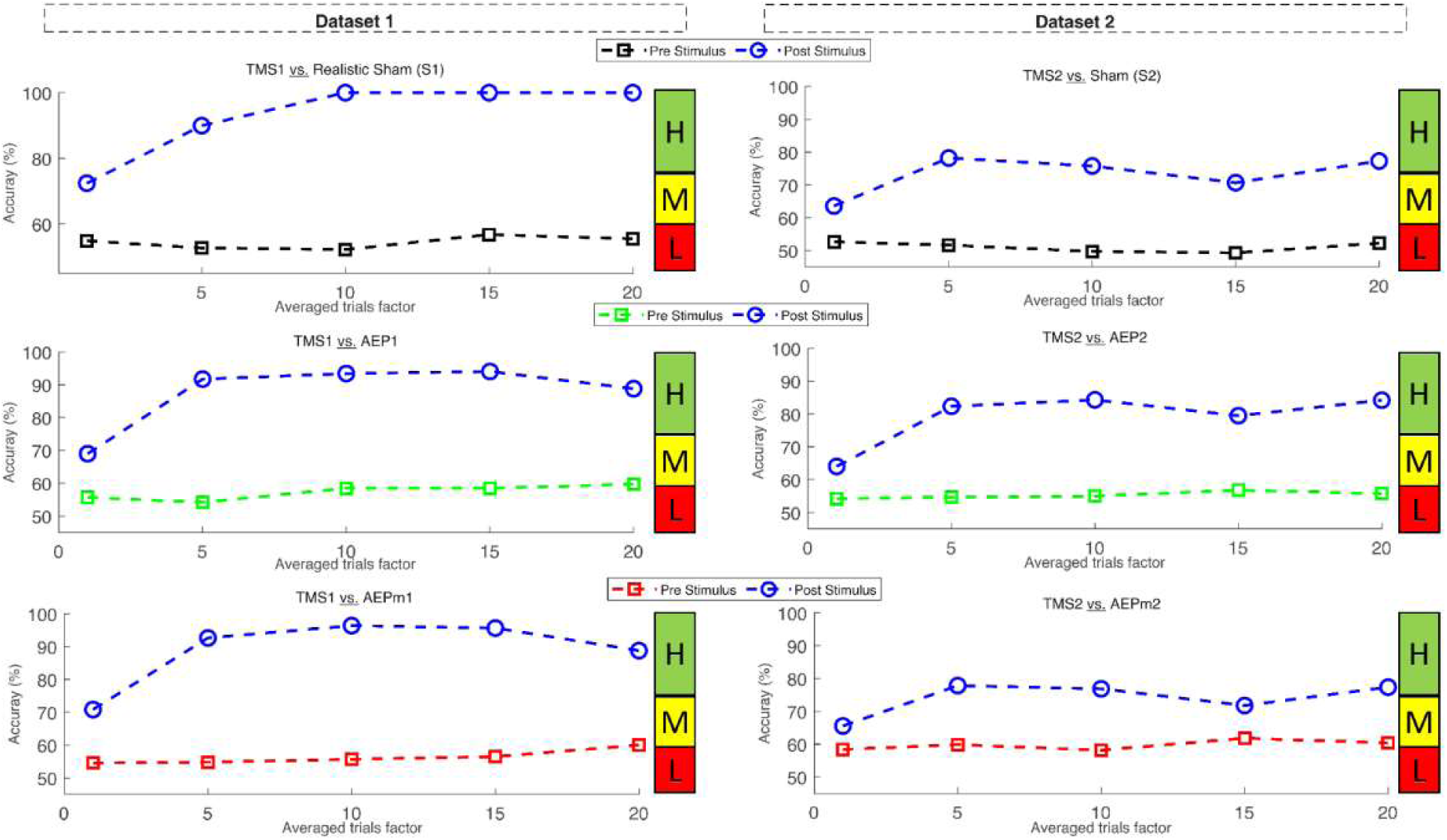
Baseline/pre-stimulus comparisons between TMS and shared sham conditions yielded low (single trial) to low/moderate (20 trials) accuracy, whereas post- stimulus comparisons showed moderate (single trial) to high (20 trials) accuracy across datasets. Accuracy rates (%, y axes) after averaging an increasing number of trials (x axes) for baseline/pre TEP vs sham comparisons (black, green and red dotted traces) and for post TMS vs. sham comparisons (blue dotted traces) for both datasets. Color bar chart shows Acc ranges: Low (L < 60%), Medium (60% ≤ M ≤ 75 %), High (H > 75 %).

**Table 2:** Main findings from between-condition comparisons of TEP and different sham stimulations. L: low Acc (<60%), M: moderate Acc (60-75%), and H: high Acc (>75%) for single trial and 20 trials Acc across subjects.

| Comparisons | Model performance |  |  |  |
| --- | --- | --- | --- | --- |
|  | Pre TMS vs. Pre sham |  | Post TMS vs. Post sham |  |
|  | Single trial<br>(Acc [%]) | Average of 20<br>trials (Acc [%]) | Single trial<br>(Acc [%]) | Average of 20<br>trials (Acc [%]) |
| **Dataset 1** |  |  |  |  |
| **TEP (TMS1) vs. Realistic sham (S1)** | L(54.87) | L(55.49) | M(72.41,) | H(100) |
| **TEP (TMS1) vs. AEP not masked (AEP1)** | L(55.62) | L(59.69) | M(68.96) | H(88.78) |
| **TEP (TMS1) vs. AEP masked (AEPm1)** | L(57.41) | M(62.79) | M(70.82) | H(88.77) |
| **TEP (TMS1) vs. High intensity Sham (HES1)** | L(50.81) | L(59.32) | M(74.25) | H(100) |

| Dataset 2 |  |  |  |  |
| --- | --- | --- | --- | --- |
| TEP (TMS2) <u>vs.</u> Sham (S2) | L(52.64) | L(52.25) | M(63.57) | H(77.27) |
| TEP (TMS2) <u>vs.</u> AEP not masked (AEP2) | L(54.15) | L(55.72) | M(64.03) | H(84.20) |
| TEP (TMS2) <u>vs.</u> AEP masked (AEPm2) | L(58.37) | M(60.36) | M(65.54) | H(77.37) |
| TEP (TMS2) <u>vs.</u> ES (LES2) | M(70.20) | M(73.56) | H(77.79) | H(93.20) |
| TEP (TMS2) <u>vs.</u> TEP not masked (TSMnm2) | L(50.20) | L(47.10) | L(57.36) | M(66.87) |

#### Post-stimulus comparisons of TMS vs. sham conditions shared across datasets yielded moderate (single trial) to high (20 trials) accuracy

In Dataset 1, moderate to high accuracy levels were obtained for post vs. post comparisons, including TMS1 vs. S1 (Figure 4, top left panel, blue dotted line, and Table 2, Acc range=[72.41%-100%]), TMS1 vs. AEP1 (Acc range=[68.96%-88.78%], Figure 4, middle left panel, blue dotted line, and Table 2), and TMS1 vs. AEPm1 (Acc range=[70.82%-88.77%], Figure 4, left bottom panel, blue dotted line, and Table 2). Similar findings were obtained in Dataset 2 (Figure 4, right panels, blue dotted lines, and Table 2).

#### TMS vs. unique sham conditions confirmed between-condition trends except for post TMS2 vs. post TMSnm2

Low/moderate accuracy rates for the pre-TMS vs. pre-sham comparisons along with moderate to high accuracy for post-TMS vs. post-sham comparisons were also observed for the sham conditions unique to each dataset (Supp. Figures 2-3 and Table 2), except for post TMS2 vs. post TMSnm2, which included TEP with and without noise masking and yielded low to moderate accuracy levels (Supp. Figure 4, Table 2).

#### Single subject and other performance parameters confirmed group accuracy findings

Single-subject between-condition comparisons confirmed the findings from group analyses for accuracy (Figure 5). Furthermore, sensitivity, specificity, AUC, and F1-score provided results comparable to those obtained with accuracy for both datasets (Supp. Figures 2-10).

**Figure 5.**
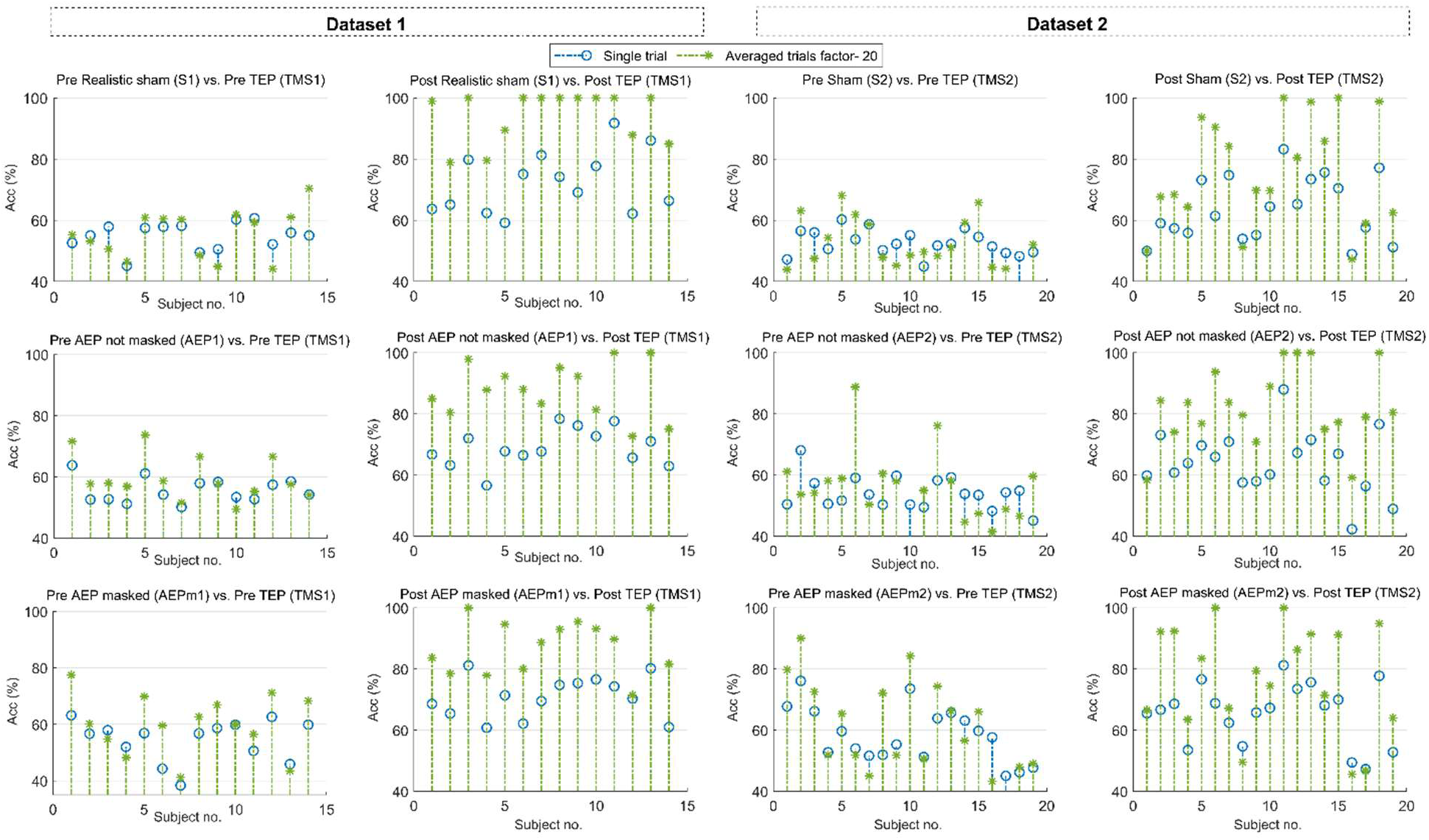
Single subject confirmed group-level findings for pre- vs. pre- and post- vs. post-comparisons between TMS and shared sham conditions across datasets. Individual accuracy rates for single trial (blue circles) and the average of 20 trials (green asterisks) of pre-TMS vs. pre-sham (first and third column) and post-TMS vs. post-sham (second and fourth column) comparisons are shown for both datasets.

## Discussion

We applied a sequence-to-sequence BiLSTM ML algorithm to identify EEG responses to TMS and several sham stimulation conditions using two different TMS-EEG datasets. Our ML approach yielded: 1) at least moderate accuracy for post- vs. pre-TMS comparisons at the single trial level; 2) high accuracy for the post- vs. pre-TMS comparisons for 20 trials, which was higher than the correspondent post- vs. pre-sham comparisons; 3) low accuracy for comparisons between pre-TMS and pre-sham conditions, for both single trial and 20 trials; 4) moderate accuracy for post-TMS vs. post-sham comparisons at single trial and high accuracy for 20 trials. These findings were consistent across datasets as well as for single subject analyses, thus indicating that TEPs can be discerned from sham responses after just a few trials and at the single subject level.

The BiLSTM achieved moderate accuracy in discriminating pre- vs post-stimulus responses for TMS1 and TMS2 after a single trial. As for traditional event-related potentials, TEPs are usually observed after averaging several trials [40–42]. An intriguing implication of single trial findings is that our ML algorithm can detect TEP features after each TMS pulse. Also, the performance of the BiLSTM increased with averaging (i.e., from one to twenty trials), yielding high accuracy rates for both datasets, as expected when collecting TEPs during a TMS-EEG session. Together, these findings indicate that TEPs characteristics can be accurately detected with BiLSTM even after averaging ≤20 trials.

Our ML algorithm reached high accuracy levels in discriminating pre- vs post-stimulus for both TMS1 and TMS2. However, while TMS1 yielded 100% accuracy, TMS2 reached ∼80% accuracy. The higher performance of the former is likely related to the between-dataset differences in experimental paradigms. TMS1 data were gathered using rt-TEP, which allows obtaining TEPs with higher SNR (TEPs peak-to-peak amplitude ≥6 µV [30]), by optimizing TMS parameters (e.g., coil position, stimulation intensity) during TMS-EEG sessions while minimizing stimulation-related artifacts (e.g., scalp muscle activation) and also checking for effectiveness of noise-masking [17, 30]. In contrast, TMS2 data were acquired without rt-TEP by employing 90% RMT [16]. Furthermore, consistent with the assumption that higher SNR in TMS1 vs. TMS2 was due to the TEPs, rather than to differences in TMS-EEG data quality acquisition, surrogate data analyses yielded chance-level accuracy for both TMS1 and TMS2.

S1 and S2 showed lower accuracy levels compared to TMS1 and TMS2 respectively (Figure 2). However, while S2 had low accuracy levels both for single trial and after averaging 20 trials, S1 yielded moderate to high accuracy levels. A possible explanation for this discrepancy is that S1, but not S2, delivered an electrical stimulation to superficial scalp muscles, and a significant activation of these muscles evoked a saliency-related multimodal response contaminating S1 [9]. Although some scalp muscle activation after TMS is unavoidable, this finding suggests that, when collecting TMS-EEG data, it is important to minimize the activation of such muscles.

Concerning the other sham conditions shared across datasets, the AEP and TEP not masked yielded high accuracy levels in discriminating post- vs pre-stimulus responses, while low accuracy was reported for AEP masked. Recent studies showed that AEP is a major source of contamination in TMS-EEG responses [10, 34–37], and there is an ongoing discussion about the best approach to mitigate its effects on the TEPs [6, 10, 12, 13, 35–37]. Here, AEP not masked generated a response that was detected with moderate to high accuracy; however, when applying a noise masking, the accuracy of BiLSTM dropped to low/chance level. This is an important confirmation that EEG responses due to the TMS click can be suppressed when noise masking is properly used [6, 12, 17].

Notably, the BiLSTM could discern with high accuracy TMS from sham conditions involving auditory and/or somatosensory stimulation in both TMS-EEG datasets. Differentiating TEPs due to direct cortical activation from other brain-evoked responses is a topic of great interest [14, 16, 18, 38]. Conde and coworkers found no significant differences in TEPs between sham and TMS sessions, leading the authors to propose that TEPs derive mostly from auditory and somatosensory responses, rather than from TMS direct neuronal cortical activation. Conversely, Gordon and colleagues found no evidence for interaction between TMS-EEG responses and sensory inputs in early TEPs (i.e., ≤100 ms) [18, 38]. Two recent TMS-EEG studies, which also controlled for TMS-related click and somatosensory scalp activation [16] could reliably differentiate TEPs from sham responses up to 300 ms after TMS [17]. Furthermore, a recent ML study showed that AEP- contaminated responses could be differentiated from TEPs [29]. Here, we demonstrated that TEPs could be discerned from several sham conditions with high accuracy. A notable exception was TMSnm2, which involved TMS delivered without the noise masking. The fact that TMS responses without noise masking were less discernable/more similar to real TMS than other sham conditions is an important finding further corroborating the notion that TMS does indeed evoke TEPs via direct cortical activation, rather than through the auditory pathway.

Along with high accuracy, group analyses showed high sensitivity (i.e., the ability to correctly identify a TMS condition), specificity (i.e., the ability to detect any sham condition), AUC (i.e., the degree of separability between TMS and sham conditions), and F1 score (i.e., the harmonic mean of precision, or % of TMS instances correctly identified, and recall, or % of TMS instances correctly classified). High scores in all these parameters indicate that BiLSTM identified TMS and sham conditions with high proficiency.

Single-subject findings largely mirrored group-level effects. Most individuals showed similar trends when comparing TMS1 and TMS2 vs. their baselines as well as vs. sham conditions. Our approach also revealed some across-subject variability in the responses, which likely reflects inter-individual differences in SNR and/or possible contamination of TEPs by sensory components. Given that this effect was detectable after just a handful of trials, our method offers a unique opportunity to characterize the quality of every TMS-EEG recording by evaluating accuracy in detecting TEPs.

Combined, these findings suggest that objective criteria for TMS-EEG studies can be developed and may contribute to ensure that TEPs are not significantly confounded by AEP and other artifacts. Our study further highlights the need for online procedures to control the effectiveness of TMS and contribution of confounders, informed by our ML approach, to help the TMS-EEG operators.

## Limitations/future directions

Future work will help address some of the limitations of the present study. First, our findings need to be replicated on larger TMS-EEG datasets to establish their robustness and generalizability. To do so, using the same experimental setup and data analysis processing is highly desirable, if not necessary. Here, employing the rt-TEP graphical user interface (GUI), individual TMS dosing [30], and appropriate auditory masking [10] yielded higher, virtually perfect accuracy. Future work should therefore explore the possibility of integrating BiLSTM into real-time GUIs, such as rt-TEP [30], to obtain high-quality TEPs that can be standardized across TMS-EEG groups. Second, the current findings should be extended beyond the motor cortex. While the motor cortex is the most investigated area with TMS-EEG, other regions, including the dorsolateral prefrontal cortex, are important, due to their implications in the neurobiology of neuropsychiatric disorders [4, 39]. Thus, future work should examine the accuracy of BiLSTM in non-motor cortical areas and clinical populations. Third, we utilized a sequence-to-sequence ML approach (i.e., LSTM-based network) [23–26, 31, 40]. This approach should be compared with other ML methods (e.g., attention-based networks, large language models [41–47]), which will allow establishing whether accuracy can be further increased and whether self-supervised and unsupervised approaches [42, 43] can differentiate TEPs from sham responses. Nonetheless, this study already established moderate to high accuracy rates between TMS and different sham conditions with LSTM. Finally, auditory and somatosensory evoked potentials most commonly affect the TEPs [9, 48–50]. Besides auditory noise masking [23–25] and other experimental/data analysis approaches to mitigate these confounds [48], future ML studies should quantify whether/how much each of these potential contaminants is present in the TEPs. As a first, important step in that direction, here we showed that TEPs after active TMS are discernible from various sham stimulations even after a handful of trials and at the single-subject level using a BiLSTM ML approach.

## Data and code availability

The data and code that support the findings of this study are available upon request.

## Acknowledgment

This work was supported by R01MH125816 NIMH grant awarded to Fabio Ferrarelli.

---

**Supplementary Figure 1:**
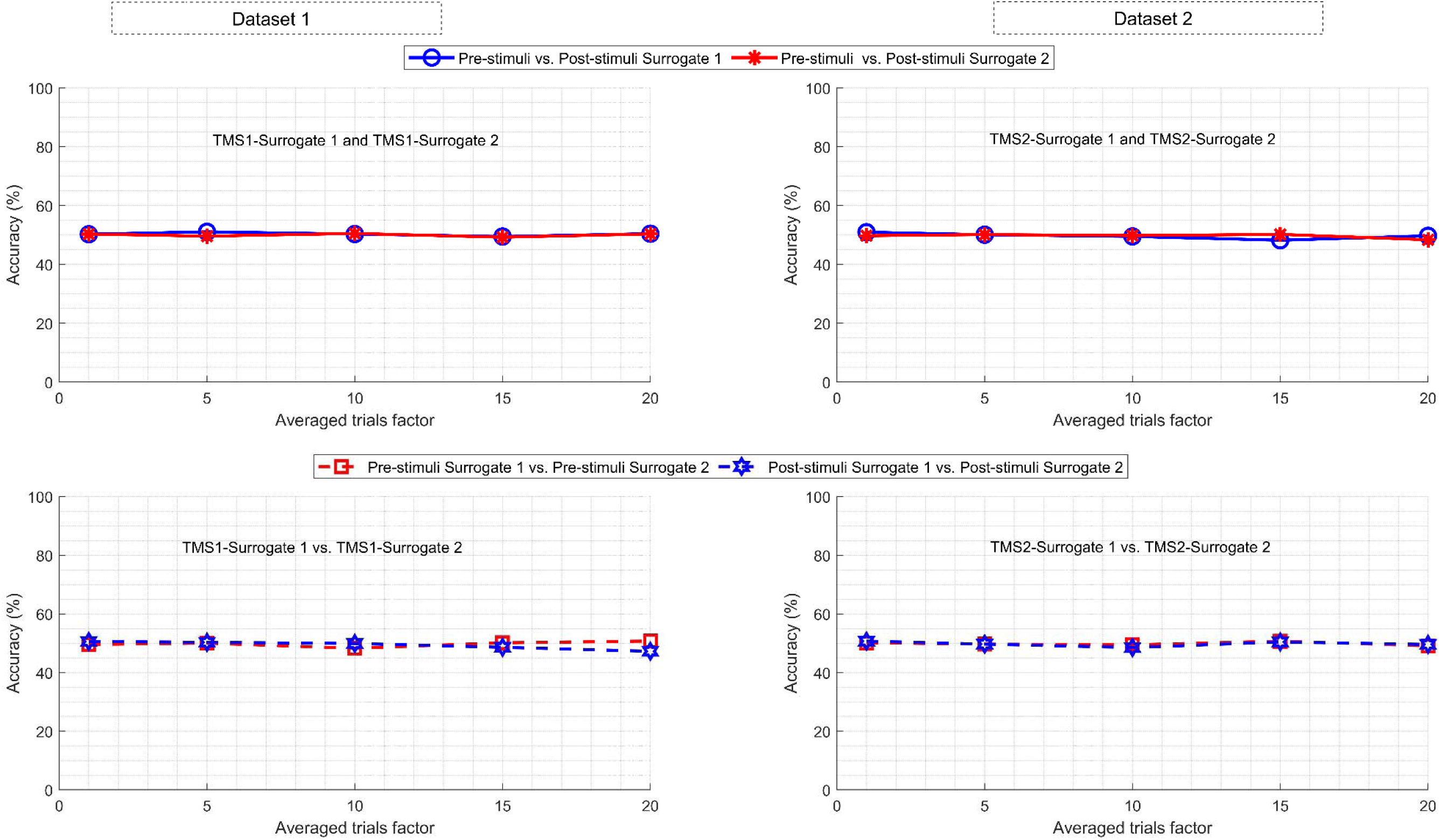
Surrogate analysis results. TMS1 and TMS2 surrogate analyses showed chance-level accuracy with both single trial and average of 20 trials analyses.

**Supplementary Figure 2:**
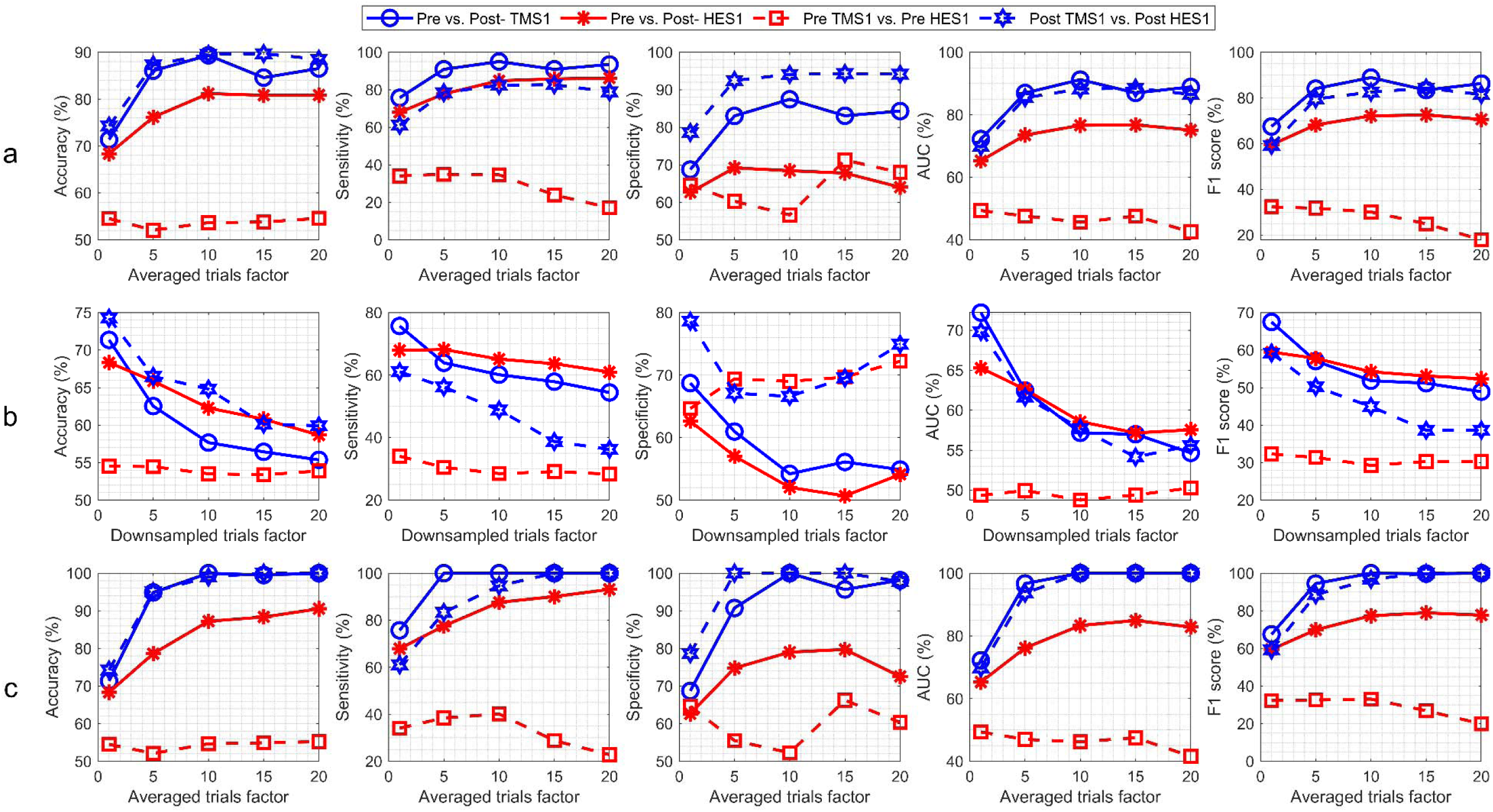
Model LOO results for TMS1vs. HES1 for Dataset 1. From left to right, accuracy, sensitivity, specificity, AUC, and F1-score are shown. Panel (a) displays the group validation results for single trial and averaged trials (i.e., n=5, etc.) in between- and within-session comparisons. Panel (b) displays the same results after accounting for each down-sampled trials factor (i.e., n=5, etc.) in between- and within-session comparisons. Panel (c) shows the adjusted values, after accounting for the reduction in the total number of trials due to the averaged trials factor (i.e., n=5, etc.).

**Supplementary Figure 3:**
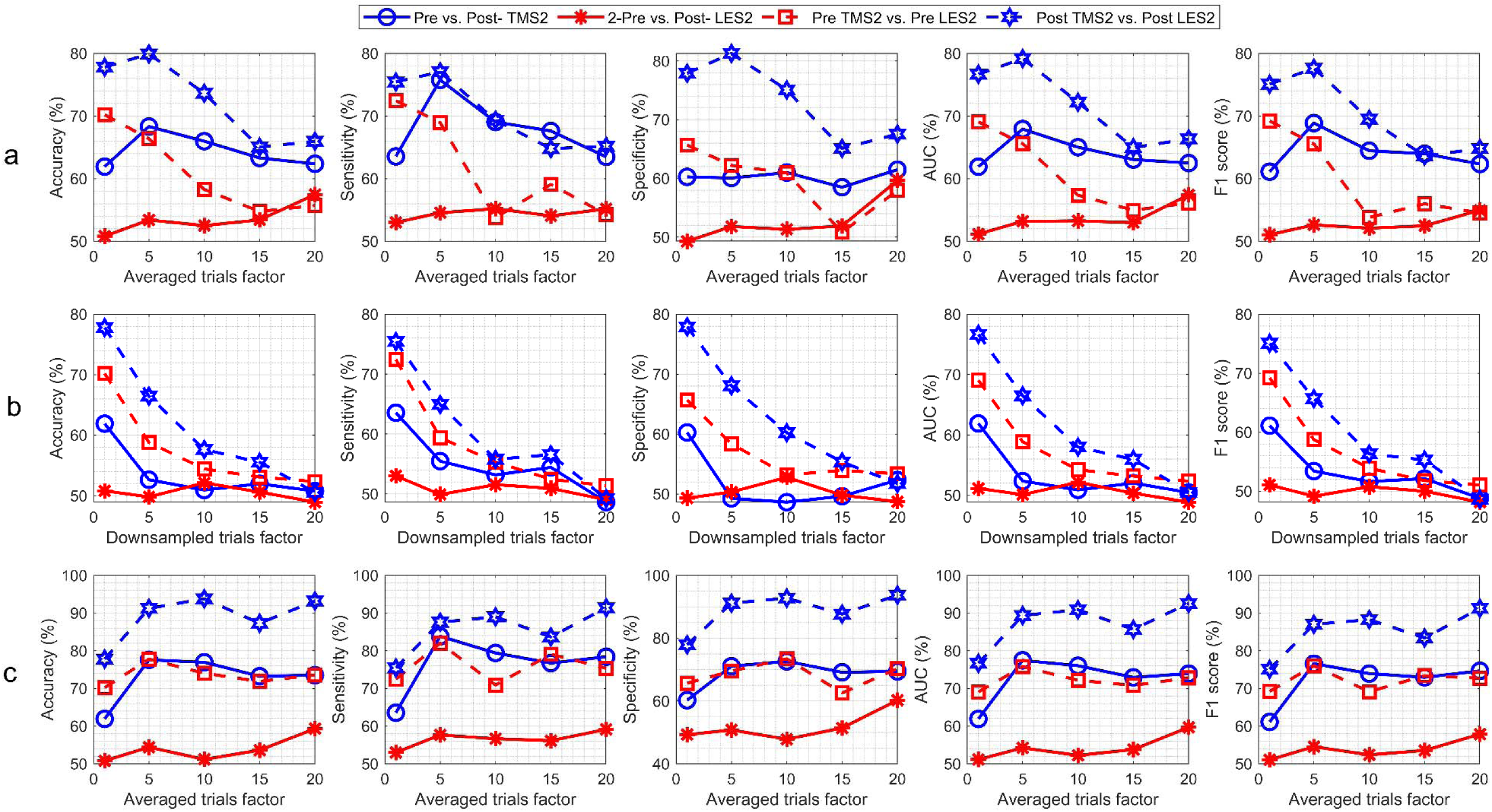
Model LOO results for single subjects for TMS2 vs. LES2 comparison for Dataset 2; the top panel shows the accuracy rates for single trial and averaged trials (i.e., n=5, etc.) in between- and within-session comparisons. The middle panel displays the same results after accounting for each down-sampled trials factor (i.e., n=5, etc.), while panel (c) shows the adjusted values after accounting for the reduction in the total number of trials due to the averaged trials factor (i.e., n=5, etc.).

**Supplementary Figure 4:**
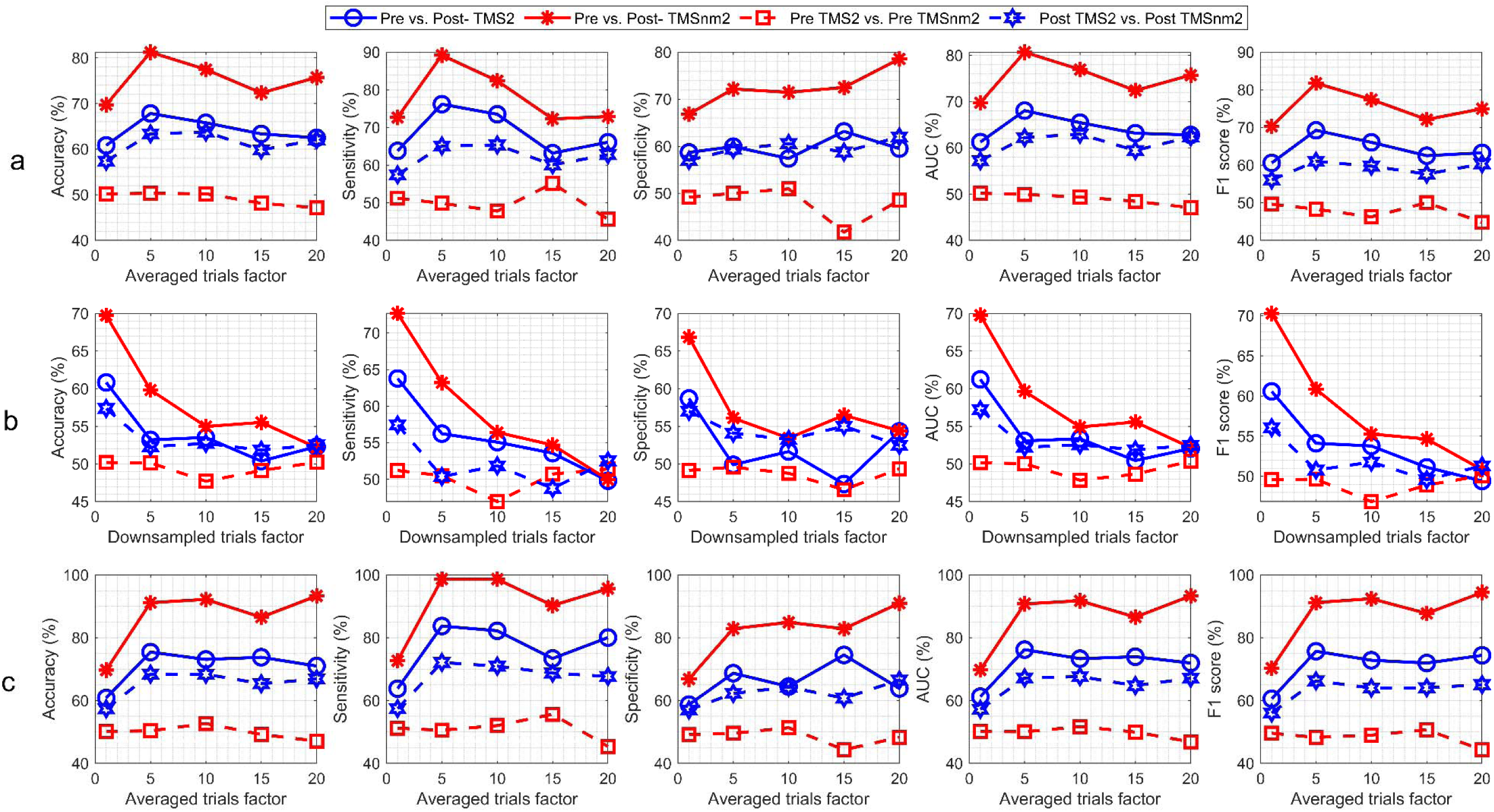
Model LOO results for TMS2vs. TMSnm2 for Dataset 2. The top panel shows the accuracy rates for single trial and averaged trials (i.e., n=5, etc.) in between- and within-session comparisons. The middle panel displays the same results after accounting for each down-sampled trials factor (i.e., n=5, etc.), while panel (c) shows the adjusted values after accounting for the reduction in the total number of trials due to the averaged trials factor (i.e., n=5, etc.).

**Supplementary Figure 5:**
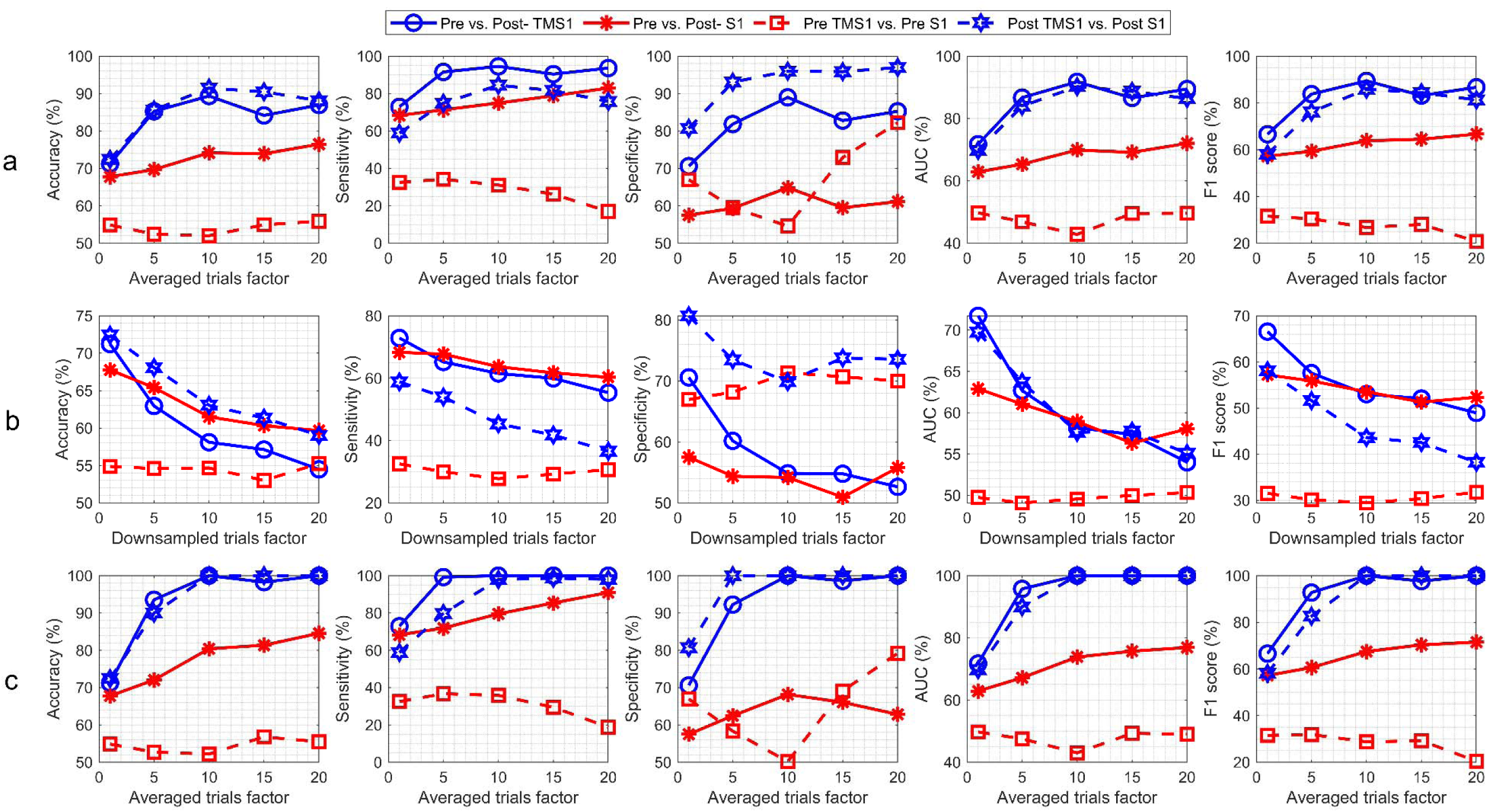
Model LOO results for TMS1vs. S1 for Dataset 1.

**Supplementary Figure 6:**
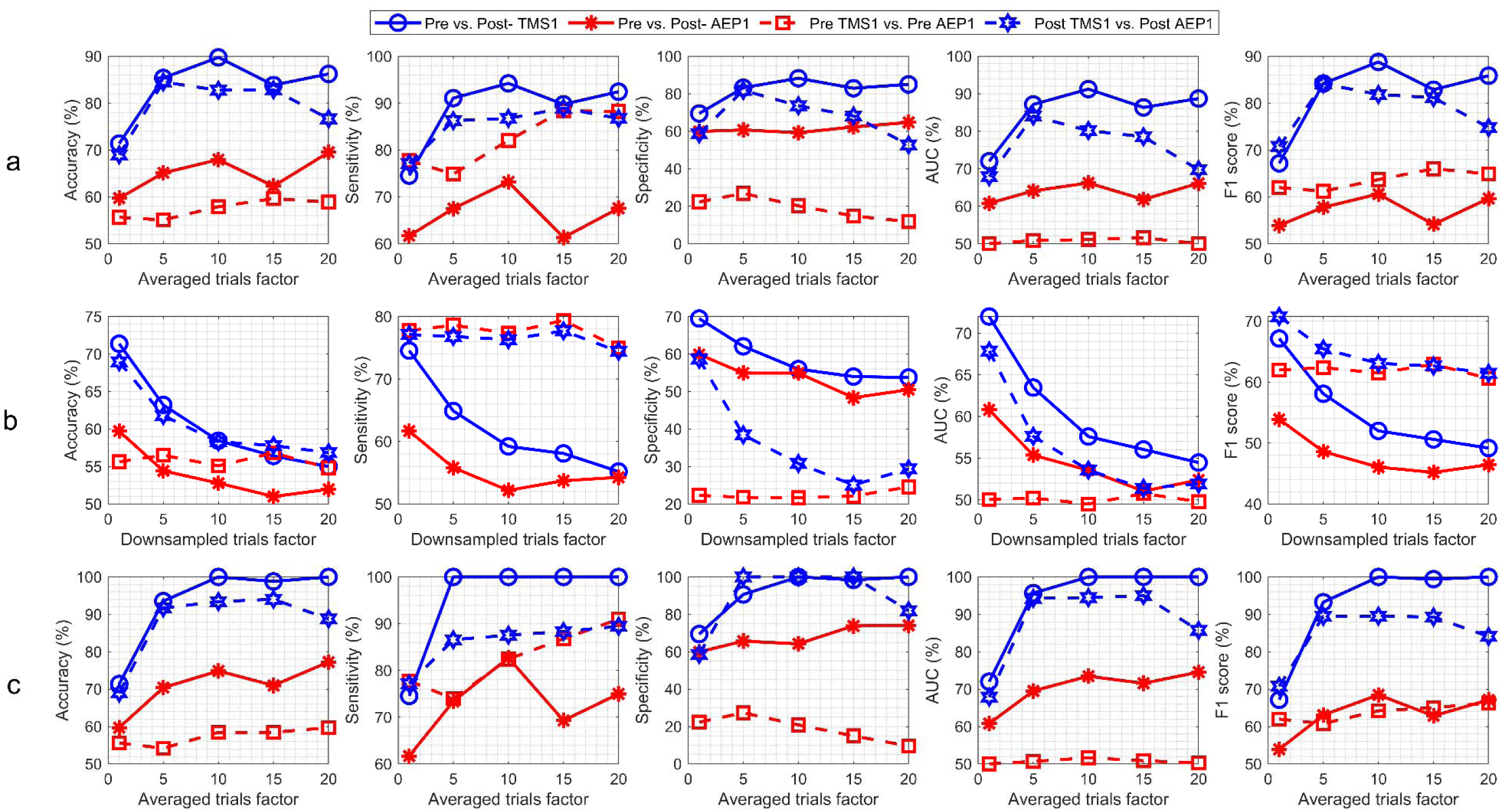
Model LOO results for TMS1 vs. AEP1 for Dataset 1.

**Supplementary Figure 7:**
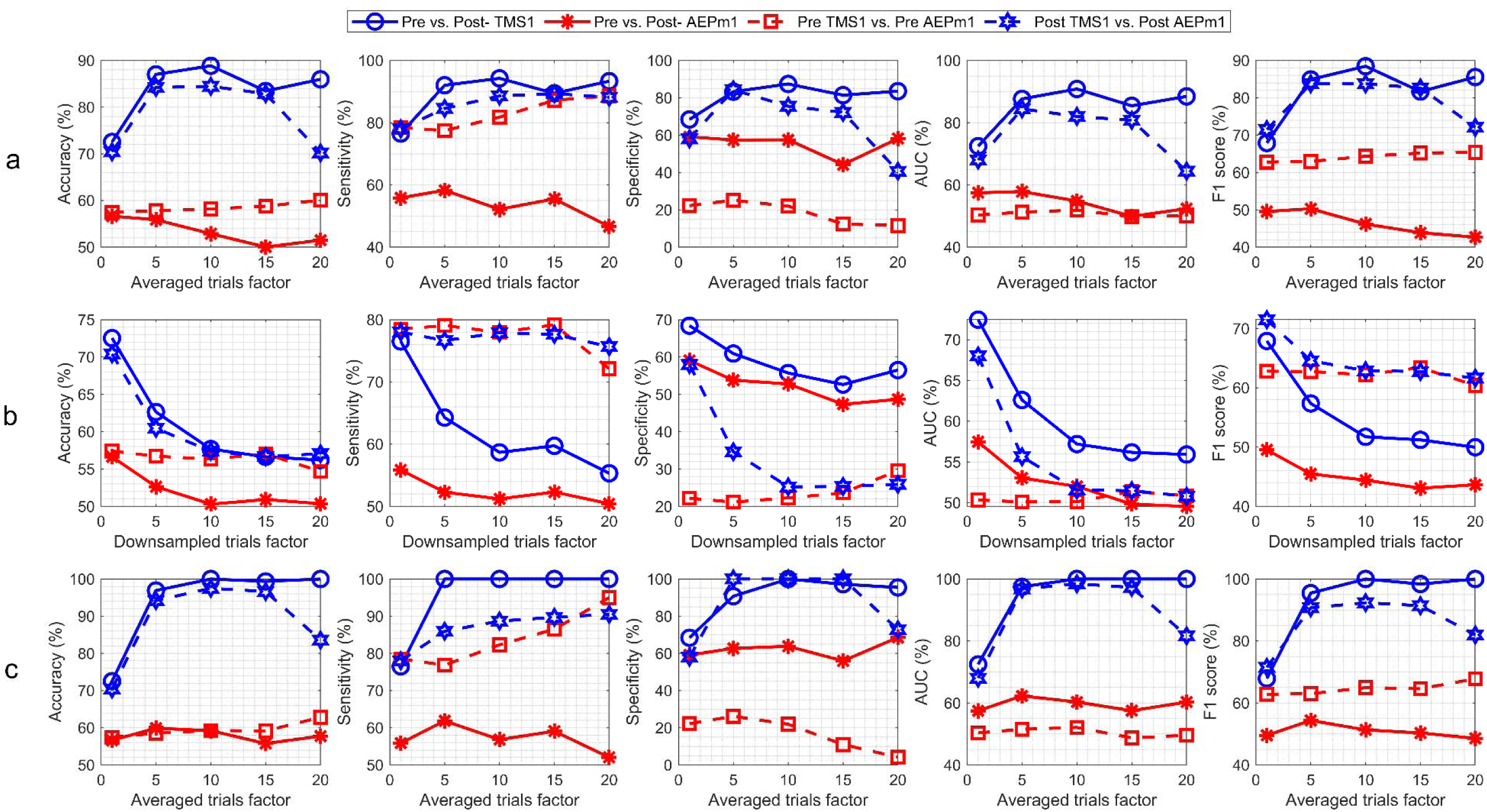
Model LOO results for TMS1 vs. AEPm1 for Dataset 1.

**Supplementary Figure 8:**
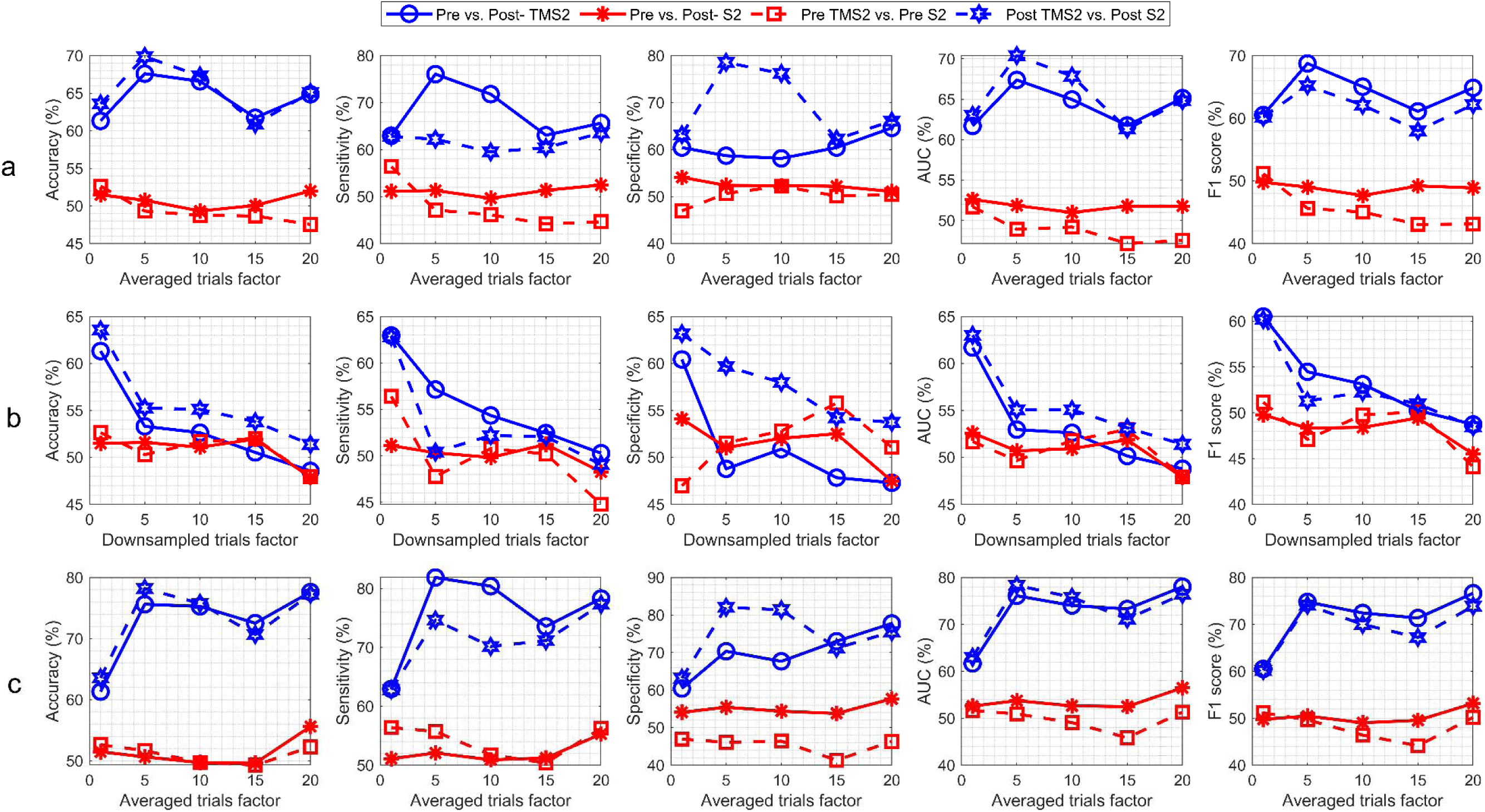
Model LOO results for TMS2 vs. S2 for Dataset 2.

**Supplementary Figure 9:**
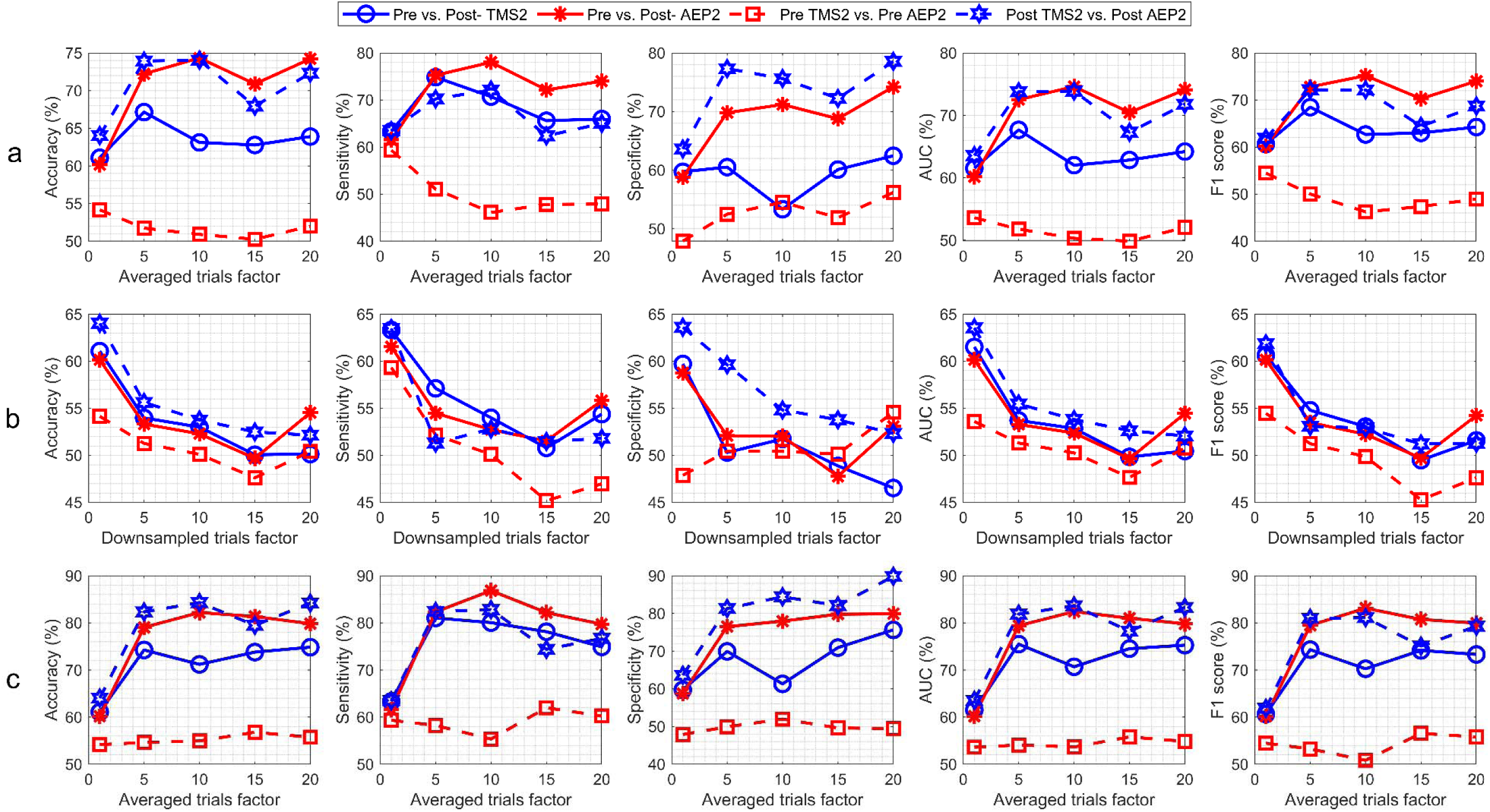
Model LOO results for TMS2vs. AEP2 for Dataset 2.

**Supplementary Figure 10:**
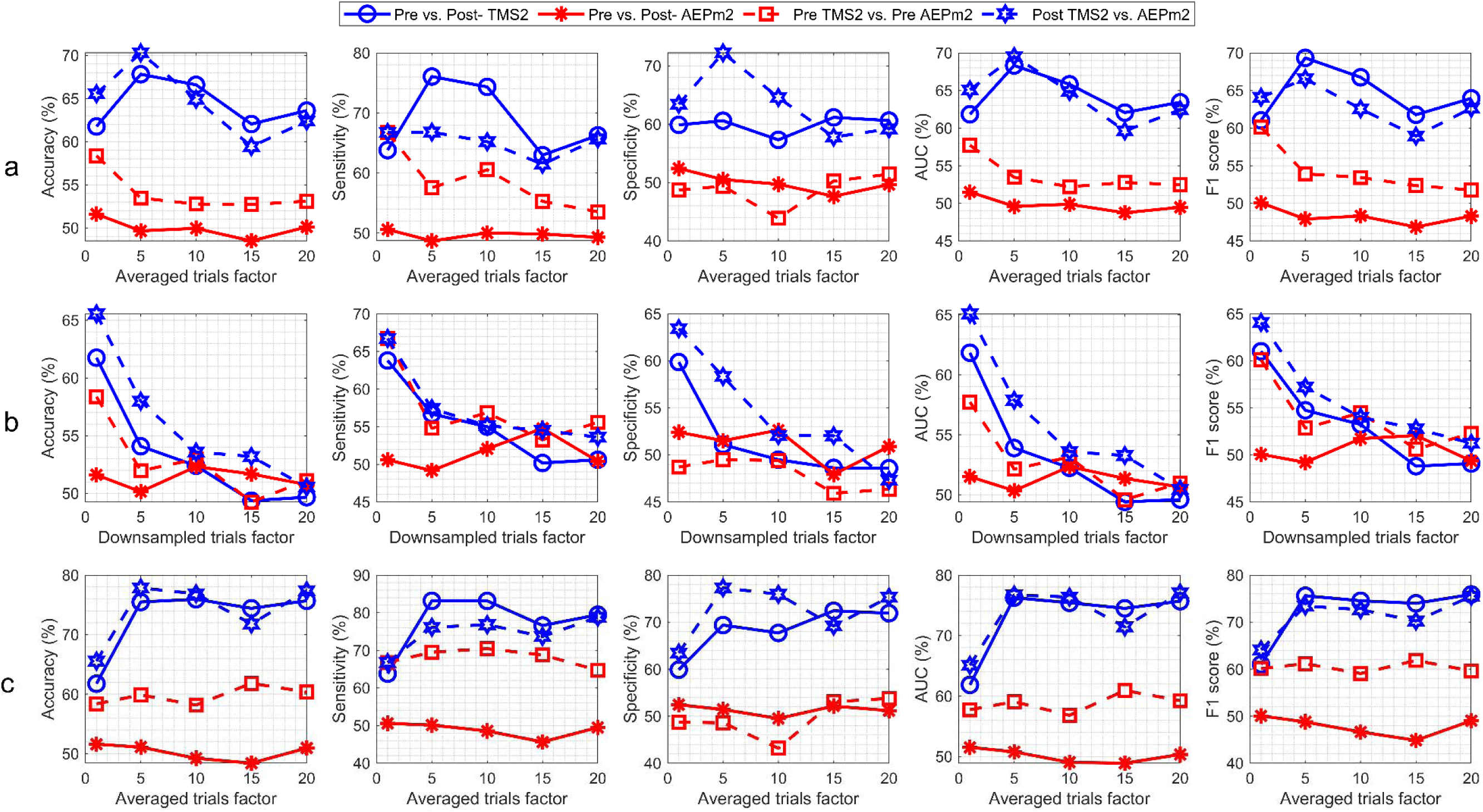
Model LOO results for TMS2 vs. AEPm2 for Dataset 2.

**Supplementary Figure 11:**
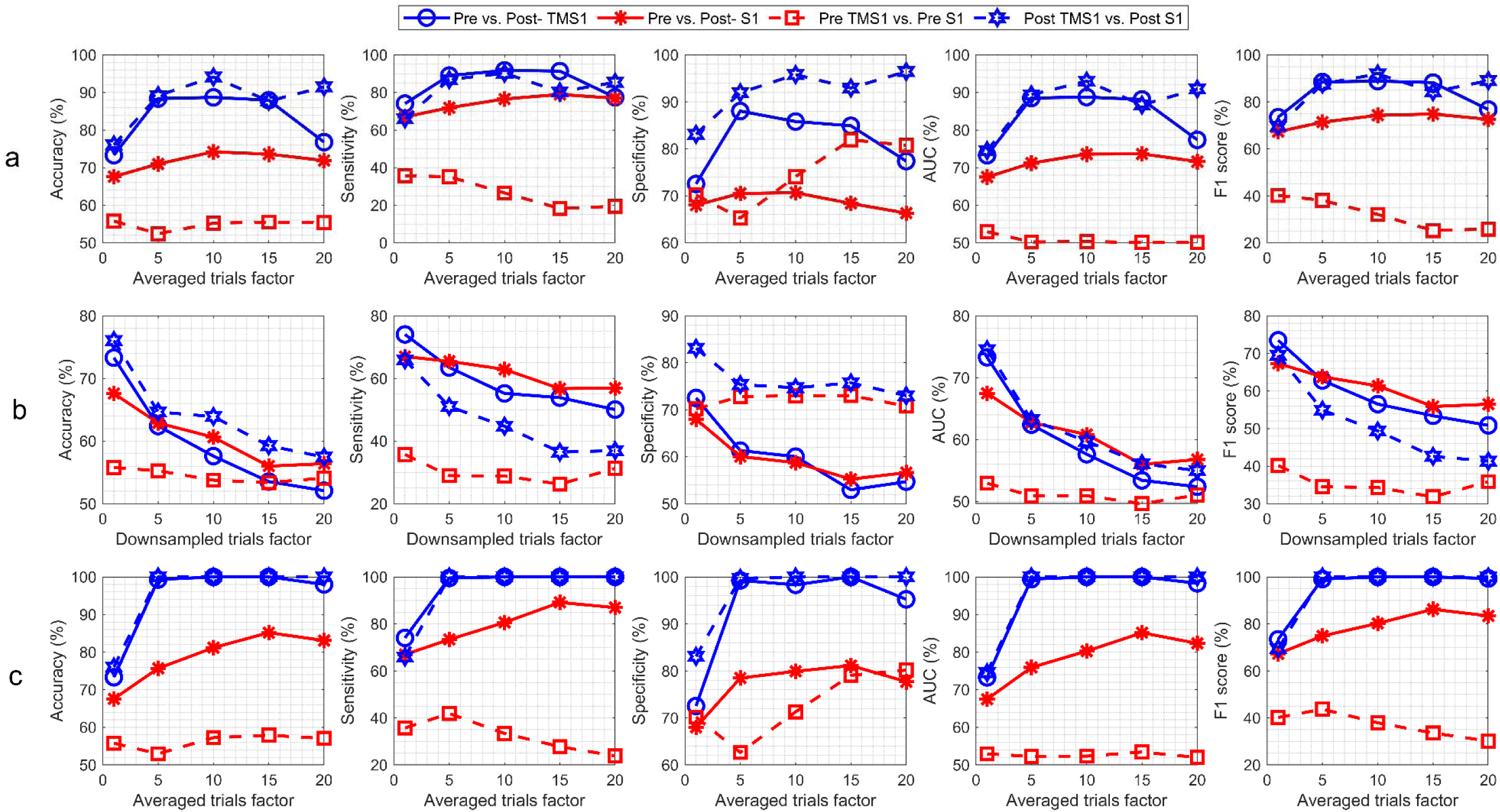
Model CV results for TMS1 vs. S1 for Dataset 1.

**Supplementary Figure 12:**
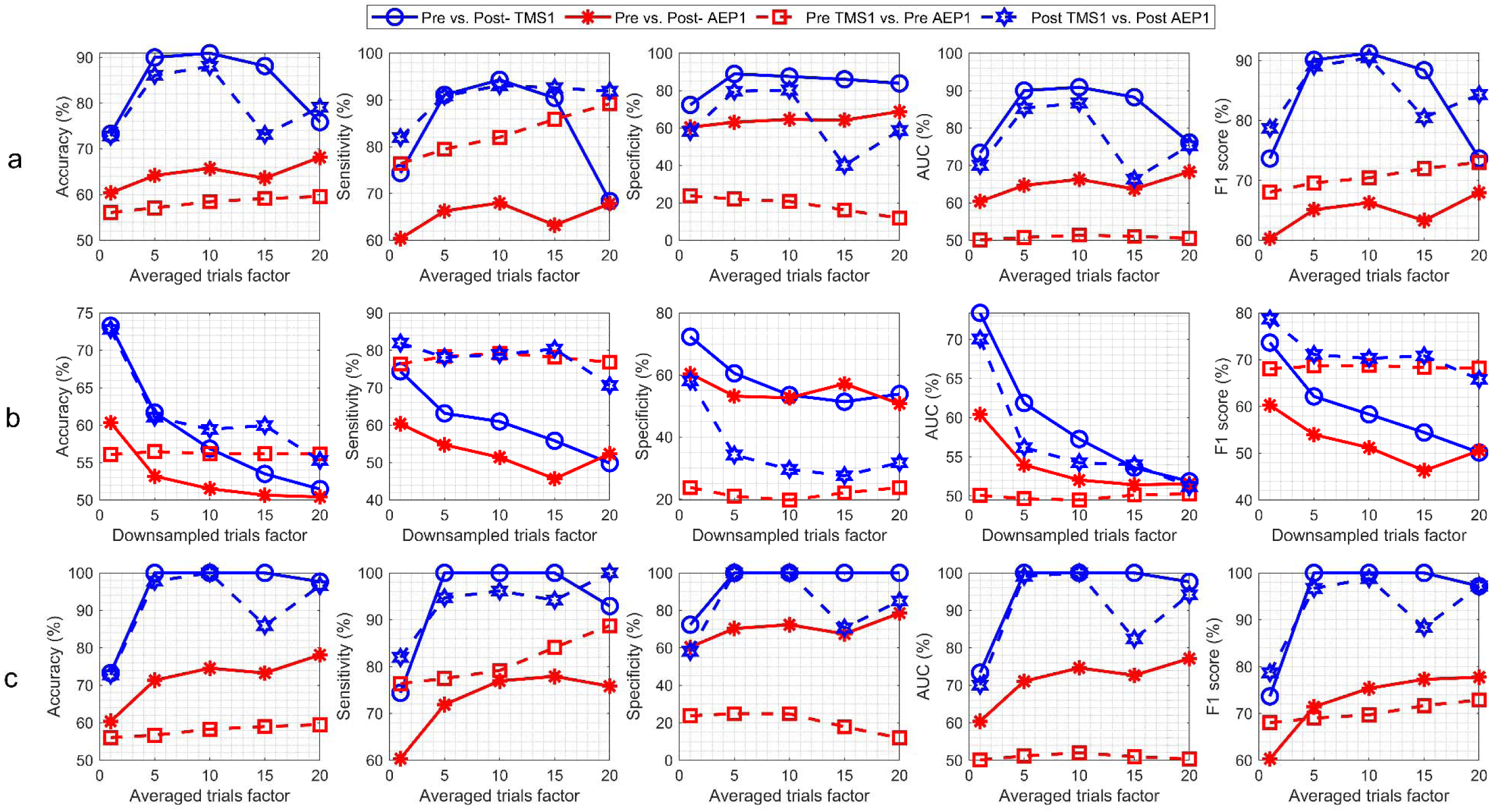
Model CV results for TMS1 vs. AEP1 for Dataset 1.

**Supplementary Figure 13:**
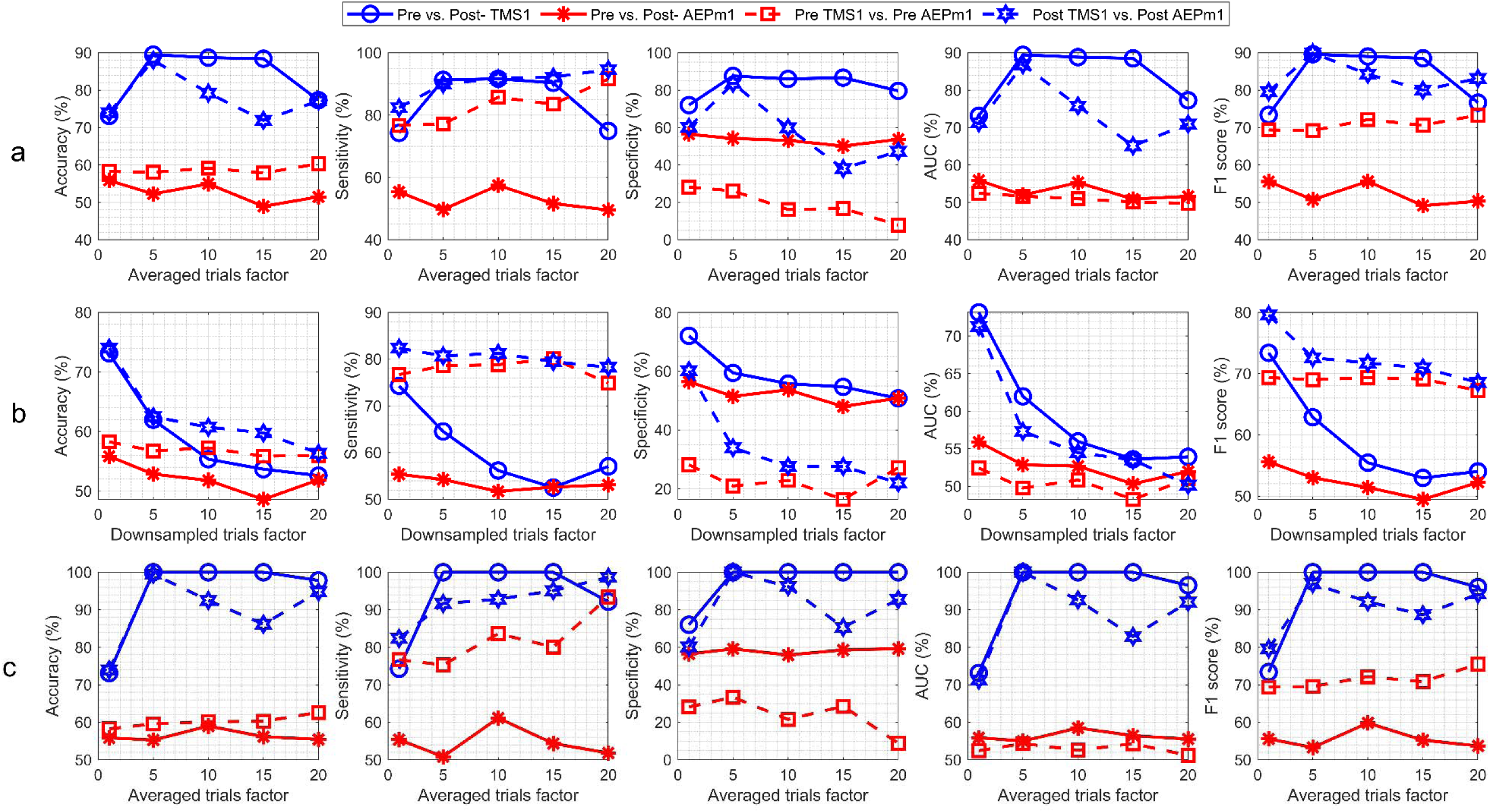
Model CV results for TMS1 vs. AEPm1 for Dataset 1.

**Supplementary Figure 14:**
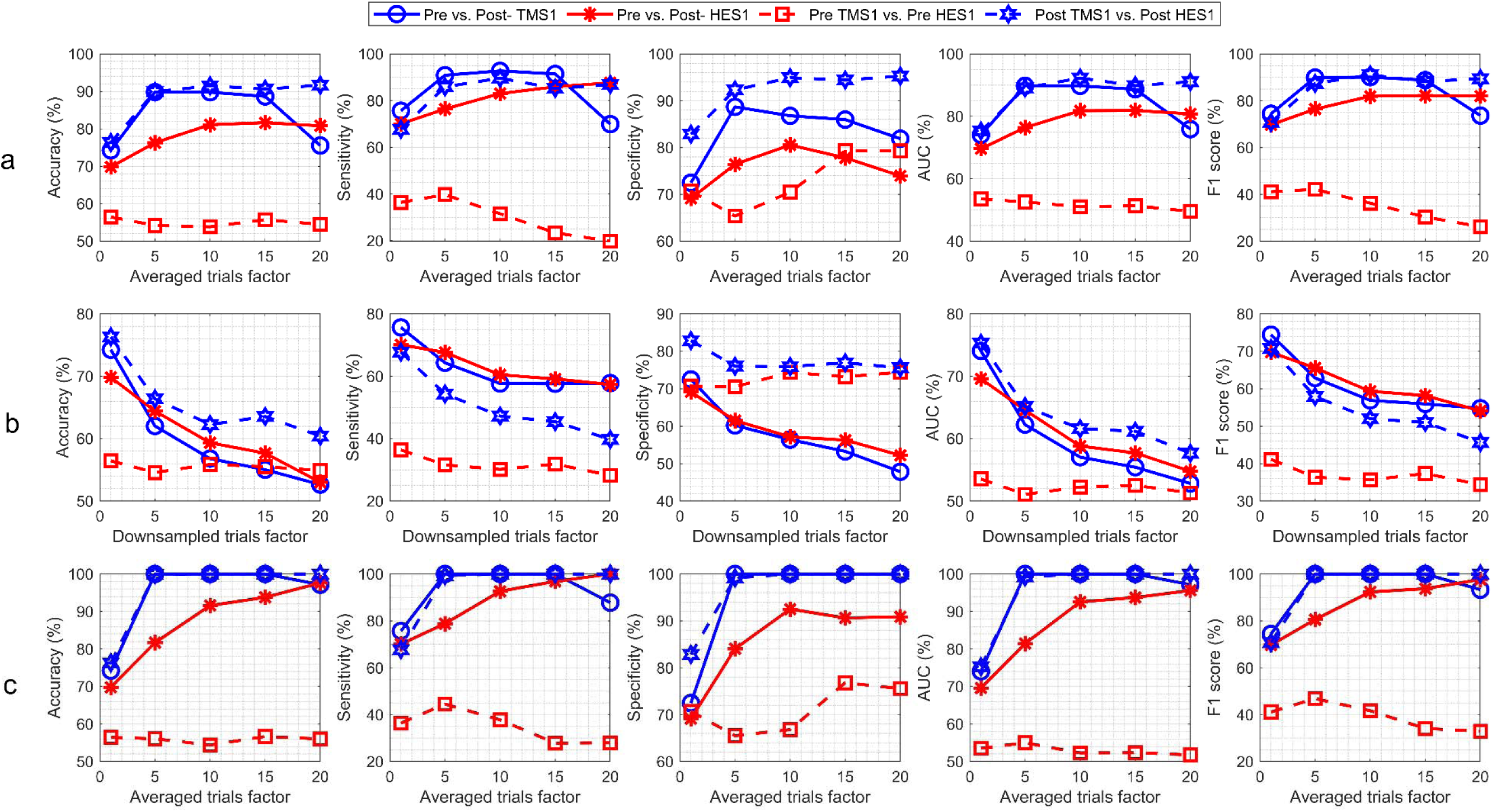
Model CV results for TMS1 vs. HES1 for Dataset 1.

**Supplementary Figure 15:**
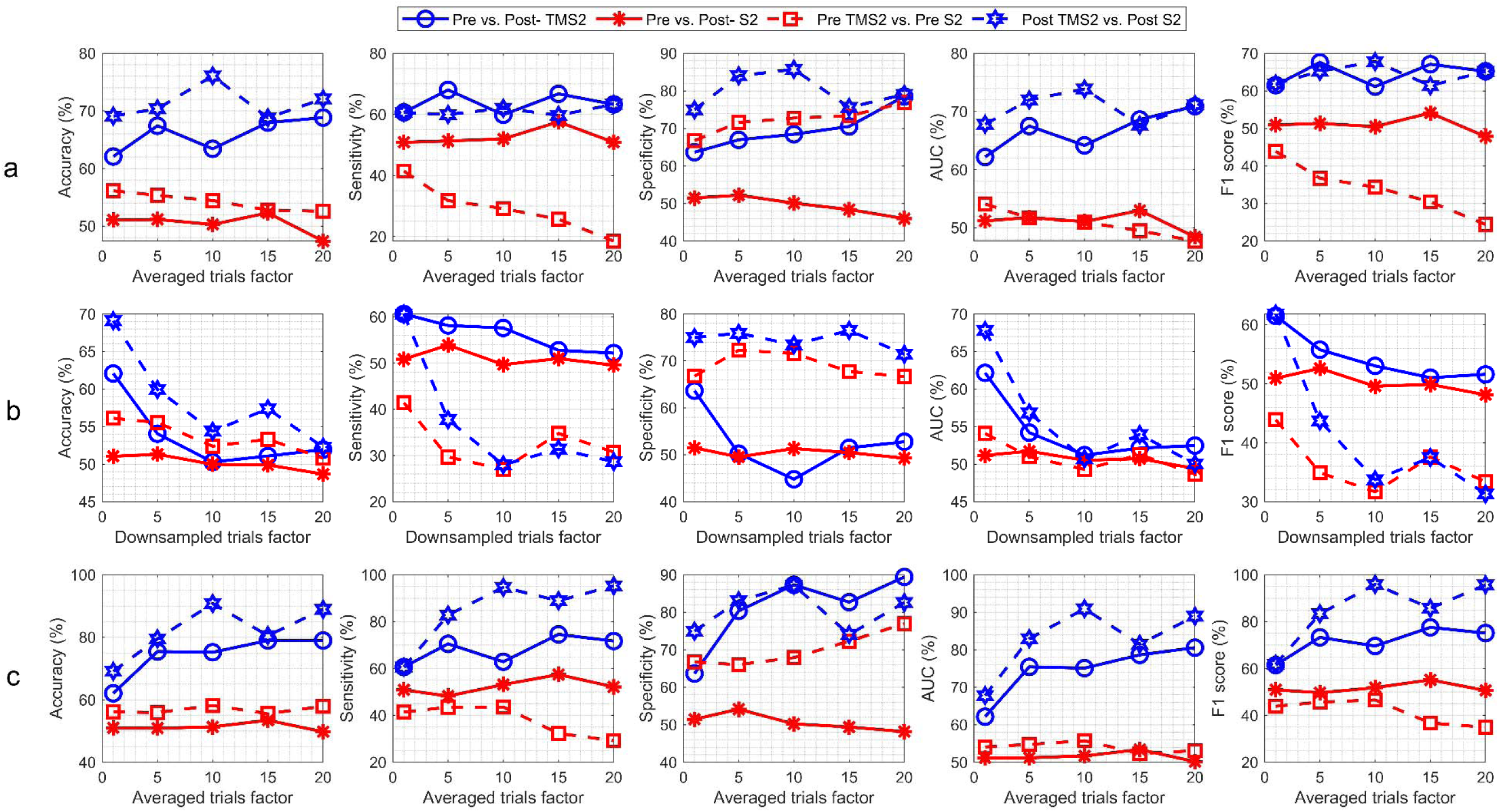
Model CV results for TMS2 vs. S2 for Dataset 2.

**Supplementary Figure 16:**
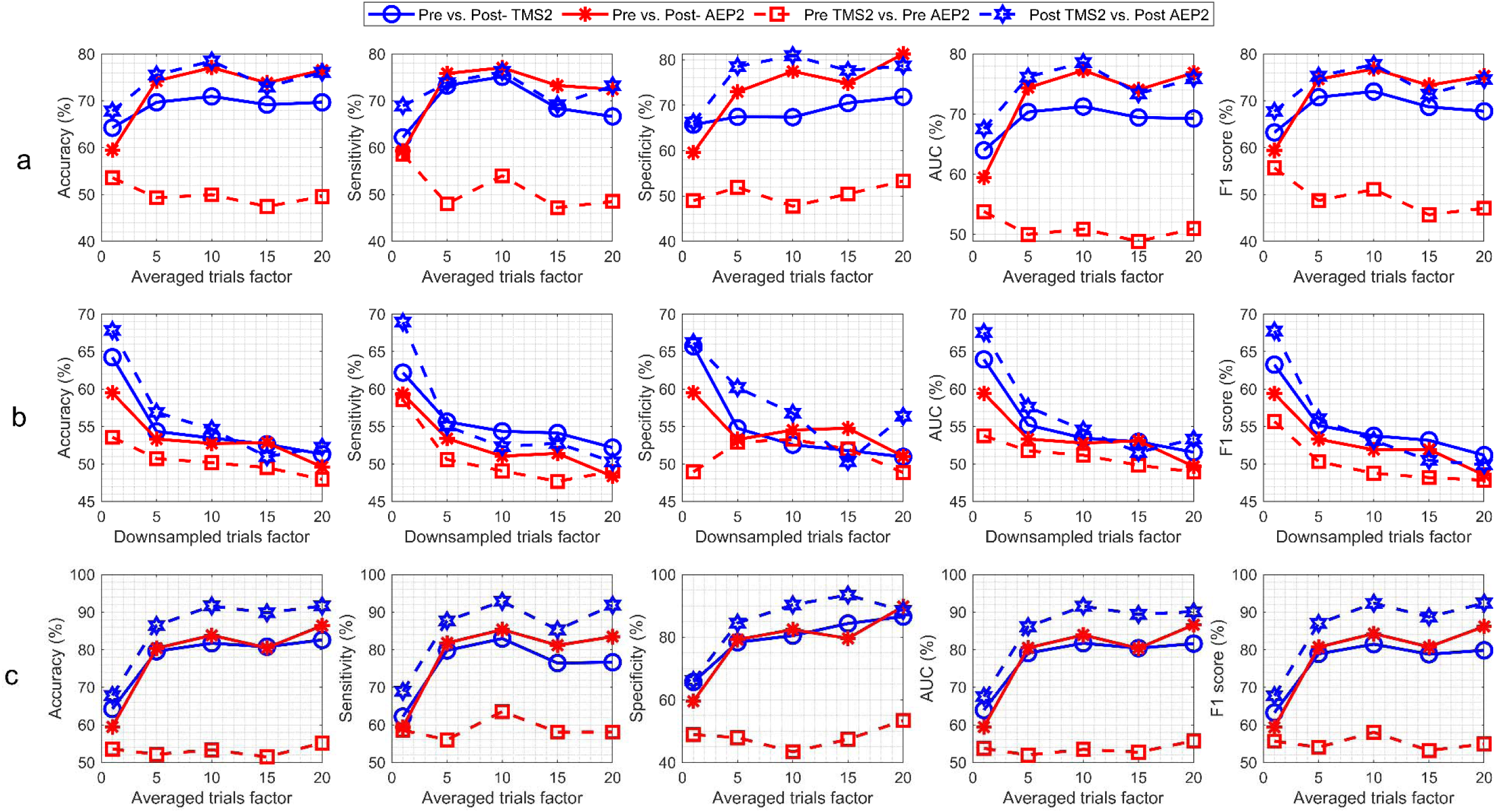
Model CV results for TMS2 vs. AEP2 for Dataset 2.

**Supplementary Figure 17:**
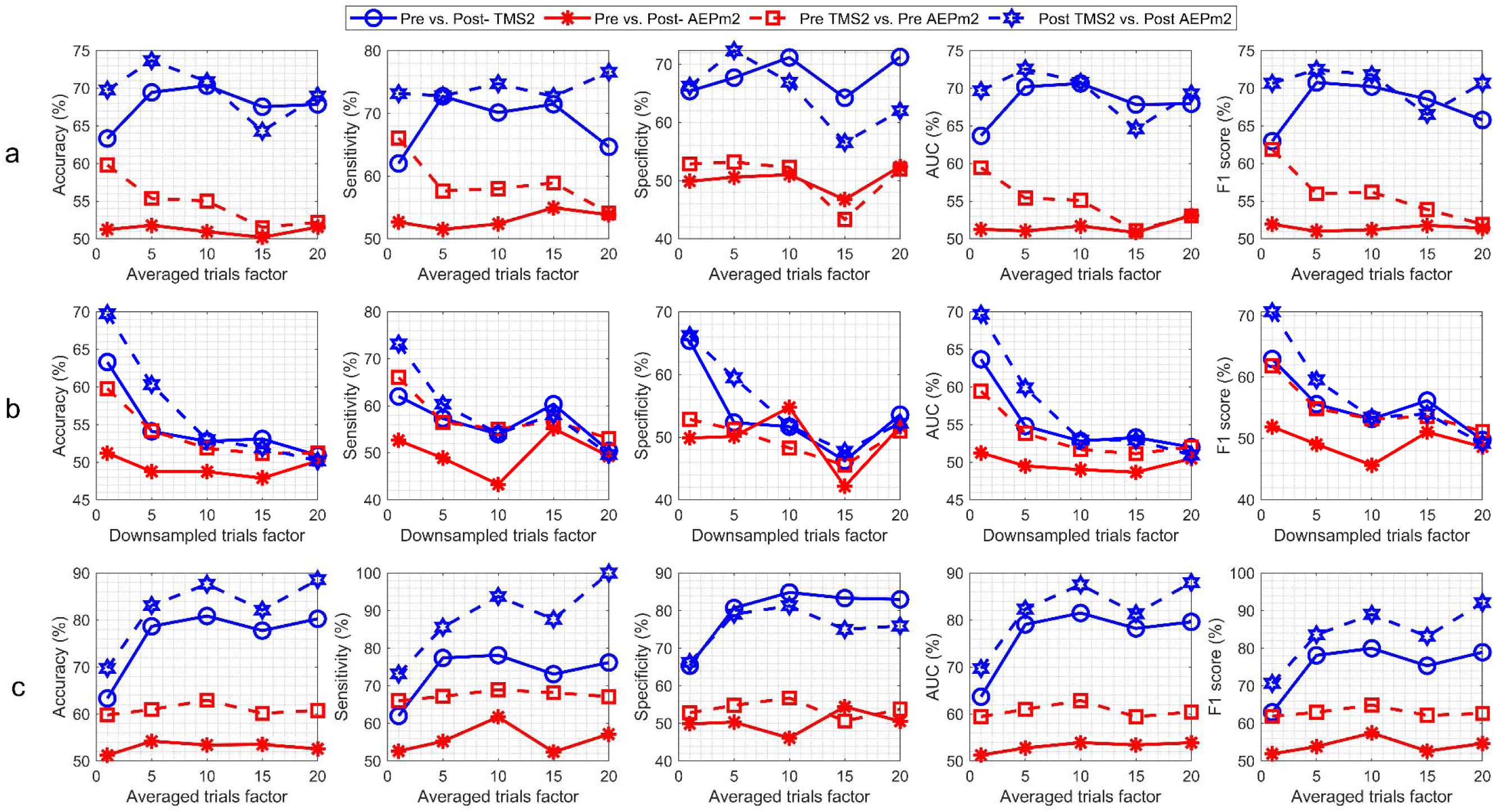
Model CV results for TMS2 vs. AEPm2 for Dataset 2.

**Supplementary Figure 18:**
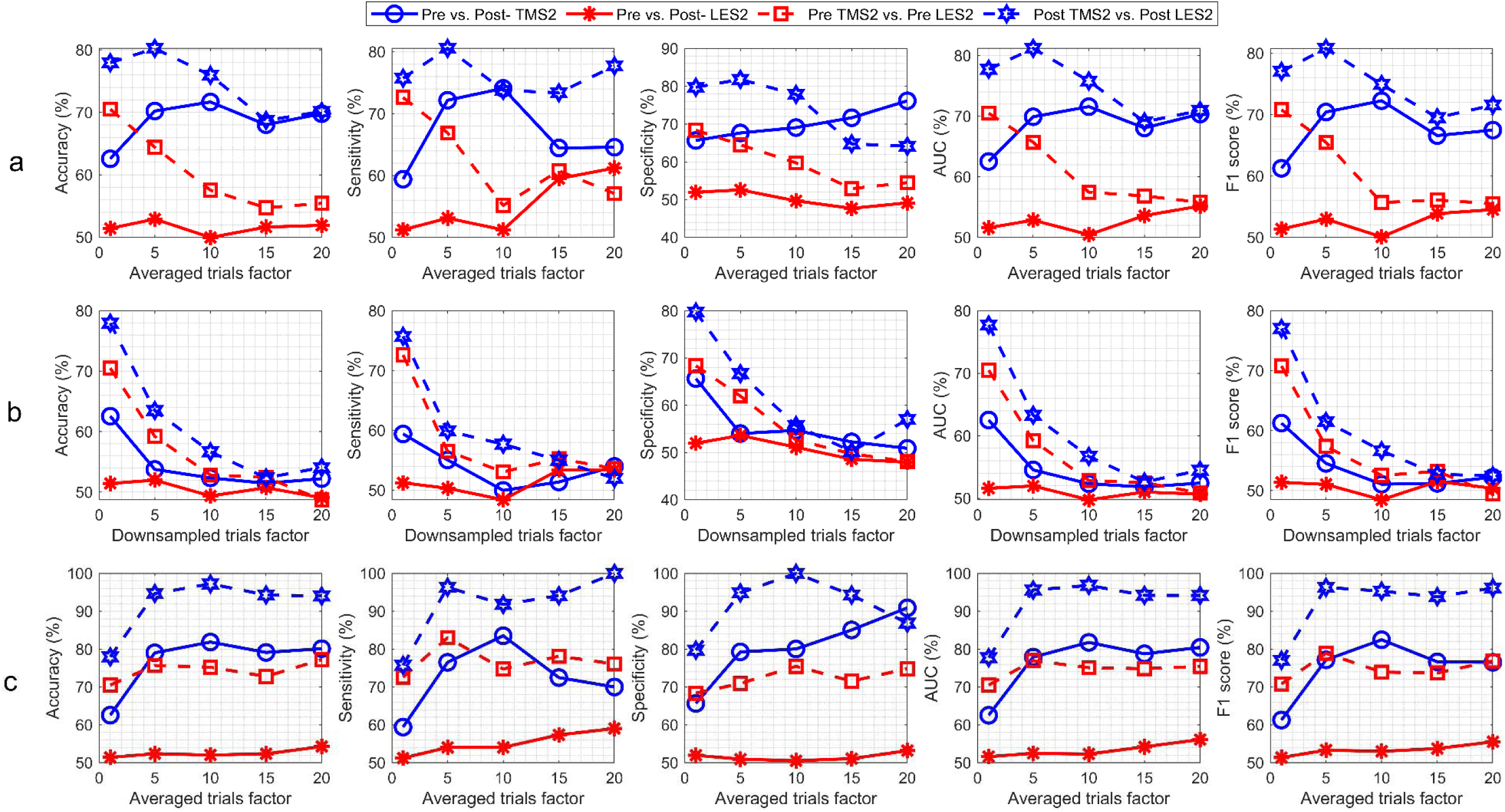
Model CV results for TMS2 vs. LES2 for Dataset 2.

**Supplementary Figure 19:**
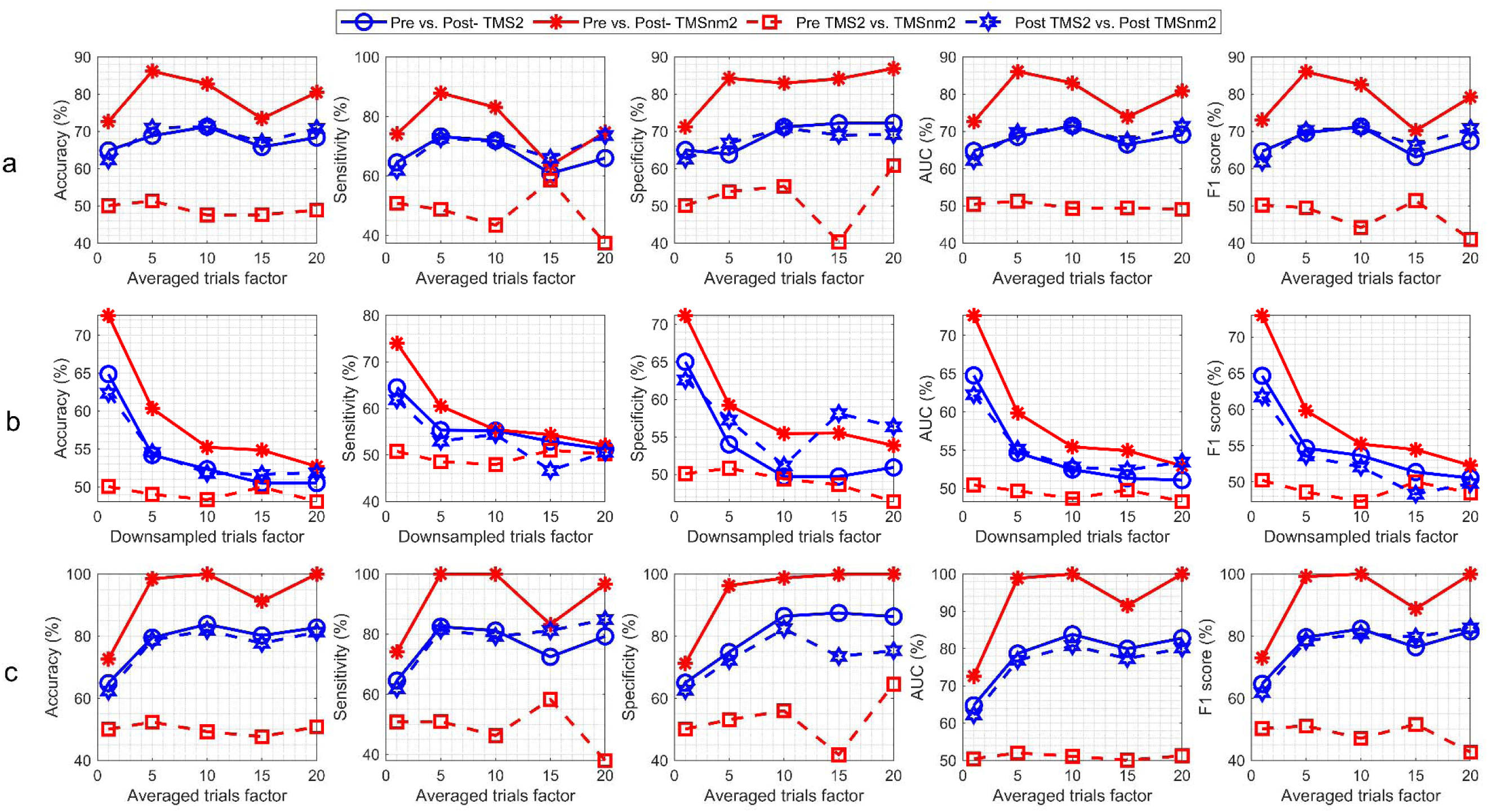
Model CV results for TMS2 vs. TMSnm2 for Dataset 2.

## References

[1] M. Bortoletto, D. Veniero, G. Thut, and C. Miniussi, “The contribution of TMS–EEG coregistration in the exploration of the human cortical connectome,” Neuroscience & Biobehavioral Reviews, vol. 49, pp. 114–124, 2015.

[2] M. Massimini, F. Ferrarelli, S. Sarasso, and G. Tononi, “Cortical mechanisms of loss of consciousness: insight from TMS/EEG studies,” Archives italiennes de biologie, vol. 150, no. 2/3, pp. 44–55, 2012.

[3] A. G. Casali et al., “A theoretically based index of consciousness independent of sensory processing and behavior,” Science translational medicine, vol. 5, no. 198, pp. 198ra105-198ra105, 2013.

[4] R. E. Kaskie and F. Ferrarelli, “Investigating the neurobiology of schizophrenia and other major psychiatric disorders with Transcranial Magnetic Stimulation,” Schizophrenia Research, vol. 192, pp. 30–38, 2018.

[5] P. Manganotti, M. Acler, S. Masiero, and A. Del Felice, “TMS-evoked N100 responses as a prognostic factor in acute stroke,” Functional neurology, vol. 30, no. 2, p. 125, 2015.

[6] J. C. Hernandez-Pavon et al., “TMS combined with EEG: Recommendations and open issues for data collection and analysis,” Brain Stimulation, 2023.

[7] D. Momi, Z. Wang, and J. D. Griffiths, “TMS-evoked responses are driven by recurrent large-scale network dynamics,” Elife, vol. 12, 2023.

[8] S. Russo et al., “Thalamic feedback shapes brain responses evoked by cortical stimulation in mice and humans,” bioRxiv, 2024.

[9] M. Mancuso et al., “Somatosensory input in the context of transcranial magnetic stimulation coupled with electroencephalography: An evidence-based overview,” Neuroscience & Biobehavioral Reviews, vol. 155, p. 105434, 2023.

[10] S. Russo, et al., “TAAC-TMS Adaptable Auditory Control: A universal tool to mask TMS clicks,” Journal of neuroscience methods, vol. 370, p. 109491, 2022.

11. J. M. Ross, M. Sarkar, and C. J. Keller, “Experimental suppression of transcranial magnetic stimulation-electroencephalography sensory potentials,” Human Brain Mapping, vol. 43, no. 17, pp. 5141–5153, 2022.

[12] P. Belardinelli et al., “Reproducibility in TMS–EEG studies: A call for data sharing, standard procedures and effective experimental control,” *Brain Stimulation: Basic*, Translational, and Clinical Research in Neuromodulation, vol. 12, no. 3, pp. 787–790, 2019.

[13] V. Conde et al., “The non-transcranial TMS-evoked potential is an inherent source of ambiguity in TMS-EEG studies,” Neuroimage, vol. 185, pp. 300–312, 2019.

[14] M. Biabani et al., “Characterising the contribution of auditory and somatosensory inputs to TMS-evoked potentials following stimulation of prefrontal, premotor, and parietal cortex,” Imaging Neuroscience, vol. 2, pp. 1–23, 2024.

[15] P. C. Gordon, D. Desideri, P. Belardinelli, C. Zrenner, and U. Ziemann, “Comparison of cortical EEG responses to realistic sham versus real TMS of human motor cortex,” Brain stimulation, vol. 11, no. 6, pp. 1322–1330, 2018.

[16] L. Rocchi et al., “Disentangling EEG responses to TMS due to cortical and peripheral activations,” Brain stimulation, vol. 14, no. 1, pp. 4–18, 2021.

[17] M. Fecchio, et al., “The specific spatiotemporal evolution of TMS-evoked potentials reflects the engagement of cortical circuits,” bioRxiv, p. 2025.06.25.661535, 2025, doi: 10.1101/2025.06.25.661535.

[18] P. C. Gordon, Y. Song, B. Jovellar, P. Belardinelli, and U. Ziemann, “No evidence for interaction between TMS-EEG responses and sensory inputs,” Brain Stimulation: Basic, Translational, and Clinical Research in Neuromodulation, vol. 16, no. 1, pp. 25–27, 2023.

[19] L. A. Gemein et al., “Machine-learning-based diagnostics of EEG pathology,” NeuroImage, vol. 220, p. 117021, 2020.

[20] M.-P. Hosseini, A. Hosseini, and K. Ahi, “A review on machine learning for EEG signal processing in bioengineering,” IEEE reviews in biomedical engineering, vol. 14, pp. 204–218, 2020.

[21] A. Keihani, A. M. Mohammadi, H. Marzbani, S. Nafissi, M. R. Haidari, and A. H. Jafari, “Sparse representation of brain signals offers effective computation of cortico-muscular coupling value to predict the task-related and non-task sEMG channels: A joint hdEEG-sEMG study,” Plos one, vol. 17, no. 7, p. e0270757, 2022.

[22] A. Keihani, S. S. Sajadi, M. Hasani, and F. Ferrarelli, “Bayesian optimization of machine learning classification of resting-state EEG microstates in schizophrenia: a proof-of-concept preliminary study based on secondary analysis,” Brain Sciences, vol. 12, no. 11, p. 1497, 2022.

[23] A. Graves, S. Fernández, and J. Schmidhuber, “Bidirectional LSTM networks for improved phoneme classification and recognition,” in International conference on artificial neural networks, 2005: Springer, pp. 799–804.

[24] M. Schuster and K. K. Paliwal, “Bidirectional recurrent neural networks,” IEEE transactions on Signal Processing, vol. 45, no. 11, pp. 2673–2681, 1997.

[25] H. Salehinejad, S. Sankar, J. Barfett, E. Colak, and S. Valaee, “Recent advances in recurrent neural networks,” arXiv preprint arXiv:1801.01078, 2017.

[26] Y. Yu, X. Si, C. Hu, and J. Zhang, “A review of recurrent neural networks: LSTM cells and network architectures,” Neural computation, vol. 31, no. 7, pp. 1235–1270, 2019.

27. [27] T. y. Mwata-Velu, J. G. Avina-Cervantes, J. M. Cruz-Duarte, H. Rostro-Gonzalez, and J. Ruiz-Pinales, “Imaginary finger movements decoding using empirical mode decomposition and a stacked BiLSTM architecture,” Mathematics, vol. 9, no. 24, p. 3297, 2021.

[28] Y. Zhang and Z. Fu, “The study of EEG Recognition of Depression on Bi-LSTM based on ERP P300,” in E3S Web of Conferences, 2020, vol. 185: EDP Sciences, p. 02007.

[29] A. Cristofari, M. De Santis, S. Lucidi, J. Rothwell, E. P. Casula, and L. Rocchi, “Machine Learning-Based Classification to Disentangle EEG Responses to TMS and Auditory Input,” Brain Sciences, vol. 13, no. 6, p. 866, 2023.

[30] S. Casarotto, et al., “The rt-TEP tool: real-time visualization of TMS-Evoked Potentials to maximize cortical activation and minimize artifacts,” Journal of Neuroscience Methods, vol. 370, p. 109486, 2022.

[31] S. Hochreiter and J. Schmidhuber, “Long short-term memory,” Neural computation, vol. 9, no. 8, pp. 1735–1780, 1997.

[32] D. P. Kingma and J. Ba, “Adam: A method for stochastic optimization,” *arXiv preprint* arXiv:1412.6980, 2014.

[33] J. Li, M. Gao, and R. D’Agostino, “Evaluating classification accuracy for modern learning approaches,” Statistics in medicine, vol. 38, no. 13, pp. 2477–2503, 2019.

[34] F. Farzan, M. Vernet, M. M. Shafi, A. Rotenberg, Z. J. Daskalakis, and A. Pascual-Leone, “Characterizing and modulating brain circuitry through transcranial magnetic stimulation combined with electroencephalography,” Frontiers in neural circuits, vol. 10, p. 73, 2016.

[35] M. Poorganji, et al., “Differentiating transcranial magnetic stimulation cortical and auditory responses via single pulse and paired pulse protocols: A TMS-EEG study,” Clinical Neurophysiology, vol. 132, no. 8, pp. 1850–1858, 2021.

[36] E. M. Ter Braack, C. C. de Vos, and M. J. van Putten, “Masking the auditory evoked potential in TMS–EEG: a comparison of various methods,” Brain topography, vol. 28, pp. 520–528, 2015.

[37] K. Pastiadis, et al., “Auditory Fine-Tuned Suppressor of TMS-Clicks (TMS-Click AFTS): A Novel, Perceptually Driven/Tuned Approach for the Reduction in AEP Artifacts in TMS-EEG Studies,” Applied Sciences, vol. 13, no. 2, p. 1047, 2023.

[38] P. C. Gordon et al., “Recording brain responses to TMS of primary motor cortex by EEG–utility of an optimized sham procedure,” Neuroimage, vol. 245, p. 118708, 2021.

[39] F. L. Donati et al., “Natural Oscillatory Frequency Slowing in the Premotor Cortex of Early-Course Schizophrenia Patients: A TMS-EEG Study,” Brain Sciences, vol. 13, no. 4, p. 534, 2023.

[40] M. Kudo, J. Toyama, and M. Shimbo, “Multidimensional curve classification using passing-through regions,” Pattern Recognition Letters, vol. 20, no. 11-13, pp. 1103–1111, 1999.

[41] X. Liu et al., “Large Language Models are Few-Shot Health Learners,” arXiv preprint arXiv:2305.15525, 2023.

[42] L. Fan, L. Li, Z. Ma, S. Lee, H. Yu, and L. Hemphill, “A bibliometric review of large language models research from 2017 to 2023,” arXiv preprint arXiv:2304.02020, 2023.

[43] E. Kotei and R. Thirunavukarasu, “A Systematic Review of Transformer-Based Pre-Trained Language Models through Self-Supervised Learning,” Information, vol. 14, no. 3, p. 187, 2023.

[44] Q. Wen, et al., “Transformers in time series: A survey,” *arXiv preprint* arXiv:2202.07125, 2022.

[45] C. Yun, S. Bhojanapalli, A. S. Rawat, S. J. Reddi, and S. Kumar, “Are transformers universal approximators of sequence-to-sequence functions?,” *arXiv preprint* arXiv:1912.10077, 2019.

[46] A. Vaswani et al., “Attention is all you need,” Advances in neural information processing systems, vol. 30, 2017.

[47] M.-T. Luong, H. Pham, and C. D. Manning, “Effective approaches to attention-based neural machine translation,” *arXiv preprint* arXiv:1508.04025, 2015.

[48] J. C. Hernandez-Pavon, D. Kugiumtzis, C. Zrenner, V. K. Kimiskidis, and J. Metsomaa, “Removing artifacts from TMS-evoked EEG: A methods review and a unifying theoretical framework,” Journal of Neuroscience Methods, vol. 376, p. 109591, 2022.

[49] X. Du et al., “N100 as a generic cortical electrophysiological marker based on decomposition of TMS-evoked potentials across five anatomic locations,” Experimental brain research, vol. 235, pp. 69–81, 2017.

[50] P. C. Gordon, Y. F. Song, D. B. Jovellar, M. Rostami, P. Belardinelli, and U. Ziemann, “Untangling TMS-EEG responses caused by TMS versus sensory input using optimized sham control and GABAergic challenge,” The Journal of Physiology, vol. 601, no. 10, pp. 1981–1998, 2023.

